# Automated behavioral analysis reveals that mice employ a bait-and-switch escape mechanism to de-escalate social conflict

**DOI:** 10.1101/2024.01.12.575321

**Authors:** Rachel S. Clein, Megan R. Warren, Joshua P. Neunuebel

## Abstract

Intraspecies aggression has profound ecological and evolutionary consequences, as recipients can suffer injuries, decreases in fitness, and become outcasts from social groups. Although animals implement diverse strategies to avoid hostile confrontations, the extent to which social influences affect escape tactics is unclear. Here, we used computational and machine-learning approaches to analyze complex behavioral interactions as mixed-sex groups of mice, *Mus musculus*, freely interacted. Mice displayed a rich repertoire of behaviors marked by changes in behavioral state, aggressive encounters, and mixed-sex interactions. A prominent behavioral sequence consistently occurred after aggressive encounters, where males in submissive states quickly approached and transiently interacted with females immediately before the aggressor engaged with the same female. The behavioral sequences were also associated with substantially fewer physical altercations. Furthermore, the male’s behavioral state and the interacting partners could be predicted by distinct features of the behavioral sequence, such as kinematics and the latency to and duration of male-female interactions. More broadly, our work revealed an ethologically relevant escape strategy influenced by the presence of females that may serve as a mechanism for de-escalating social conflict and preventing consequential reductions in fitness.

## Introduction

Social animals navigate complex environments by evaluating sensory cues, assessing possible risks, integrating new information with pre-existing knowledge, and executing context-appropriate behavior (1, 2). The expression of such behavioral flexibility is essential for physiological fitness, causing animals to evolve cognitive mechanisms for responding to social cues and environmental changes (3). For example, animals use transitive inference to deduce the social rank of group members by observing the behavior of others. This information allows animals to alter their behavior based on relative hierarchal rank and perceived threats (4), thus facilitating adaptable behavioral responses.

Animals respond defensively to the presence of threats with learned and innate escape behaviors (5, 6) and features of the environment strongly influence an animal’s choice in behavioral escape strategies (7). For example, pairing a neutral environmental context with a noxious stimulus produces learned freezing behaviors (8), but predator cues or hostile interactions with conspecifics elicit natural, non-conditioned responses (9). Rodents assume a defensive posture and cautiously scan the environment when encountering predator odors or freeze in response to looming shadows (10, 11). Likewise, aggressive conspecifics prompt various escape behaviors that depend upon the proximity to the threat and the likelihood of evasion (12, 13). Prior experience also plays a role in behavioral escape strategies, as animals constantly subjected to conflicts may deliberately avoid social encounters to reduce the likelihood of future attacks (14, 15). Whether pain, predators, or aggression triggers the response, successfully executing escape strategies is vital for survival. When an animal fails to implement an appropriate escape strategy, impactful consequences such as injury or, in extreme cases, death, may occur, leading to a reduction in fitness (6). Consequently, understanding the behavioral strategies animals use to escape dangerous situations is paramount.

Systematically quantifying and evaluating the effectiveness of naturalistic escape behaviors elicited by a hostile interaction represents a formidable task. To achieve this goal, we must overcome the challenges of unbiasedly extracting and assessing discrete events underlying the diverse behavioral repertoires of individual animals. By overcoming these impediments, a more comprehensive understanding of the dynamics underlying escape and the role behavioral state plays in this evolutionarily conserved behavior is achievable. Here, we monitored the behavior of multiple, freely interacting mice enclosed in a large arena and then harnessed the power of multiple computational approaches to extract the behavior of individuals. Using aggressor-aggressed behavioral states as a centralized framework, we found a robust phenomenon in which males subjected to agonistic interactions escape costly encounters and avoid conflict by exploiting nearby females to divert the attention of the pursuing aggressor. The results highlight the sophisticated social dynamics during naturalistic behavior, demonstrate that prior social experience and behavioral state influences subsequent behavior on short timescales, and reveal a previously unidentified, novel mechanism animals use to escape hostile encounters with aggressive males.

## Results

### Quantifying Dynamic Social Behavior

To explore group dynamics, we recorded naturalistic social interaction in mixed-sex groups (n = 11, 2 males, 2 females per group) of adult mice for 5 hours (**Fig 1A-C, Methods**). An automated tracking program (16) allowed unbiased quantification of movement (n = 44 animals, median total movement = 1343.7 cm, IQR = 349.3). Each mouse roamed throughout the large arena and explored the majority of the enclosure (**Fig 1C-D, S1 Fig**). To assess exploration, we separated the arena into evenly spaced bins of three different sizes (9 cm^2^, 36 cm^2^, and 81 cm^2^). For each animal, we then quantified the percentage of bins they explored throughout the experiment. At the smallest bin size, every mouse explored over 80% of the arena. For the largest bin size, more than 96% of the cage was explored by each mouse, indicating that most of the enclosure was explored by all the mice.

**Fig. 1.**
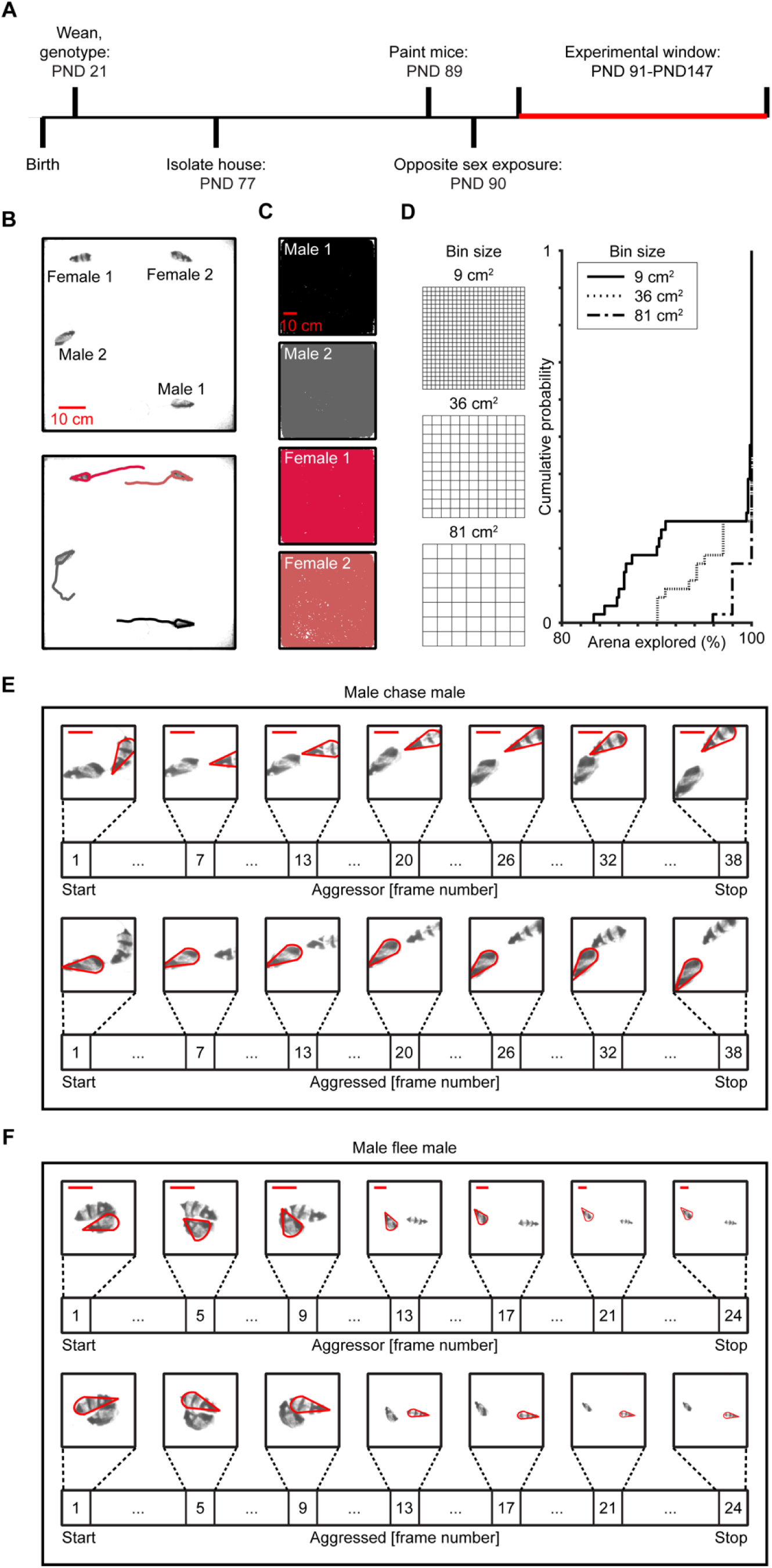
Machine learning based approaches for tracking individuals and identifying behaviors. (A) Experimental timeline. PND, post-natal day. (B) Individual mice differentiated by distinct fur patterns were tracked in a large arena (width = 76.2 cm, length = 76.2 cm, heigh = 61 cm) using automated software. Trajectories show 1 second of movement. (C) Trajectory plots showing that mice explore the arena during a 5-hour behavioral recording. (D) Cumulative probably plots quantifying the percentage of the arena explored by the mice and a schematic indicating the three binning sizes used for assessment. (E) Chase exemplar. Aggressor males were chasing (top panel), while aggressed males were being chased (bottom panel). The scale bar represents 10 cm. (F) Flee exemplar. Aggressor males were being fled from (top panel), while aggressed males fled (bottom panel). The scale bar represents 10 cm.

Because of the diversity in behavioral repertories, variation in behavior over time, and the length of each recording (17), a supervised machine-based learning program (18), which classifies user-defined behaviors, was used to identify innate agonistic behaviors (**Fig 1E-F, Methods**). Agonistic behaviors were considered events in which a male fled with (chase) or without (flee) being pursued by another male, as higher-ranked males often chase lower-ranked males and submissive males tend to flee from more dominant social partners (19). Importantly, these behaviors capture two independent behavioral states (aggressor vs. aggressed) where each mouse has a distinct, discernable role during the behavior.

Across all recordings, every male acted aggressively toward a rival (**Fig 2A-D, S2 Fig**). We observed 3,413 agonistic interactions between males (n = 22, median = 216, IQR = 331.8). Within each recording, one of the males was always more aggressive than the other male. For each male, aggregate aggression levels were calculated by comparing the total number of aggressive behaviors performed by an individual. For each recording, we quantified an aggression score where we found the difference between the aggregate aggression levels. Scores of 1 would indicate that male 1 engaged in all the aggressive behaviors, while scores of -1 would indicate that male 2 was the sole aggressor. Aggression scores ranged from -0.97 to 0.61 (**Fig 2A**). There were significant differences between the most and least aggressive males, as measured by a higher frequency (**Fig 2B**; Wilcoxon Signed Rank Test: W = 66, p = 0.001) and longer duration (**Fig 2C**; Wilcoxon Signed Rank Test: W = 62, p = 0.01) of aggressive behaviors. We observed similar patterns when partitioning aggressive behaviors into specific types (Wilcoxon Signed Rank Test; chases: W = 58, p < 0.05; flees: W = 52.5, p = 0.09). Chasing and flight occurred 2,562 and 851 times, respectively (flight median = 61, IQR = 51.3; chasing median = 174, IQR = 242.3). We also characterized the time an animal spent in a submissive state based on the time and role in an aggressive encounter (**Fig 2D**), which allowed us to establish temporal patterns of aggression for both males and categorically labeled animals as dominant and subordinate similar to prior work (20).

**Fig. 2.**
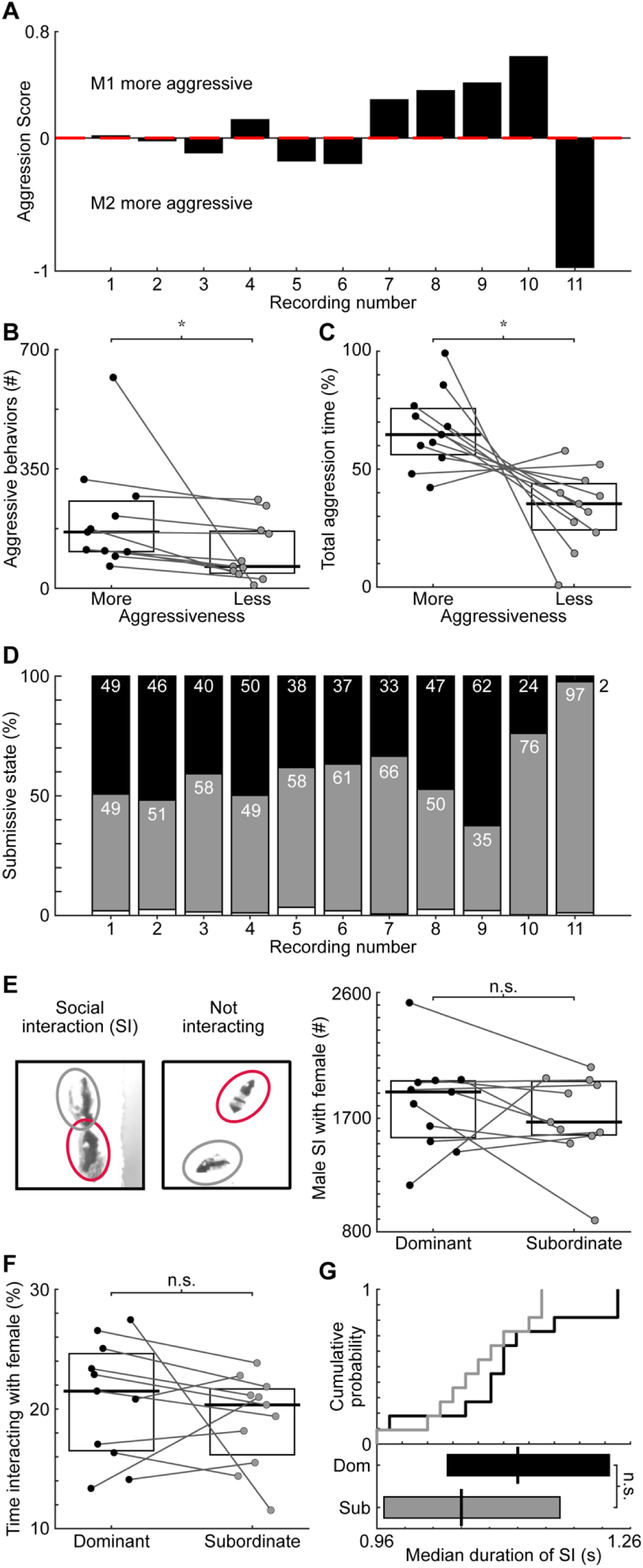
Quantifying dynamic social relationships. (A) Aggression scores show which mouse acted as the aggressor in most aggressive encounters. Recordings sorted from smallest to largest score disparities. (B) Number of aggressive behaviors per mouse. Lines connect co-recorded mice. The horizontal bars and boxes show the medians and interquartile ranges (25-75%). (C) Percentage of total aggressive time co-recorded animals spent behaving as the aggressor. (D) Overall time spent in a submissive state. Black denotes the more aggressive male, while gray denotes the less aggressive male. After quantifying the aggregate aggression levels of mice, we categorically labeled animals that spent more and less time acting aggressively as the dominant and subordinate male. (E) Left: Schematic of social interaction with females. Right: Number of social interactions (SI) between males and females. (F) Time spent interacting with females. (G) The median duration of social interactions for the dominant and subordinate animals. Top: median duration of social interactions for each animal. Bottom: median across groups. Ω = p < 0.05, * = p < 0.01, n.s. = p ≥ 0.05

In many species, females prefer dominant males, and acts of aggression serve to attract potential mates (21). Consequently, we predicted that females would socialize more with dominant males. To determine whether dominant or subordinate males interacted more with females, we identified periods where pairs of mice socialized (**Fig 2E, Supplemental Fig 2, Methods**). Social interactions involved two mice of the opposite sex spending at least 0.2 seconds within 3 centimeters of each other. Males and females frequently interacted (total interactions = 37,725, median = 3,477, IQR = 742). Contrary to expectations, the frequency (Wilcoxon Signed Rank Test, W = 131, p = 0.79), time (Wilcoxon Signed Rank Test, W = 141, p = 0.36), and duration (Wilcoxon Signed Rank Test, W = 141, p = 0.35) of female interactions with males behaving more dominantly or submissively were indistinguishable (**Fig 2E-G**).

### Behavioral State Directly Influences Male-Female Interactions

Although both dominant and subordinate males interact with females, behavioral decisions are predominately associated with an animal’s internal state (22). Thus, the purpose of social interactions may differ depending on an animal’s behavioral state. As such, the interactions between an aggressed male and female may serve as a refuge from hostile male-male interactions. To investigate this possibility, we first analyzed the temporal relationship between aggressive behaviors and opposite-sex interactions (**Fig 3A, Methods**). Across all recordings, there were 2,082 sequences where an opposite-sex social interaction followed an aggressive interaction between males. The median latency between aggressive behavior and social interaction was 1.17 seconds (IQR = 4.77 seconds). In 463 sequences, the socialization began before the aggressive behavior ended (shortest latency = -7.13 seconds, median = -0.23 seconds, IQR = 0.6 seconds). The majority (62%) of subsequent male-female interactions involved aggressed males (**Fig 3B**; aggressed state: n = 11 recordings, median = 95, IQR = 80.25; aggressive state: n = 11 recordings, median = 52, IQR = 45; Wilcoxon Signed Rank Test, W = 0, p < 0.001). The delay between hostile male-male interactions and subsequent female interactions was significantly shorter for aggressed males (**Fig 3C**; aggressive state: median = 3.15 seconds, IQR = 1.85; aggressed state: median = 0.73 seconds, IQR =1.17; Wilcoxon Signed Rank Test, W = 66, p < 0.001). Additionally, post-aggression interactions were shorter when the aggressed male engaged with the female compared to the aggressor (**Fig 3D**; aggressive state: median = 1.06 seconds, IQR = 0.35 seconds; aggressed state: median = 0.81, IQR =0.35; Wilcoxon Signed Rank Test, W = 57.50, p = 0.03).

**Fig. 3.**
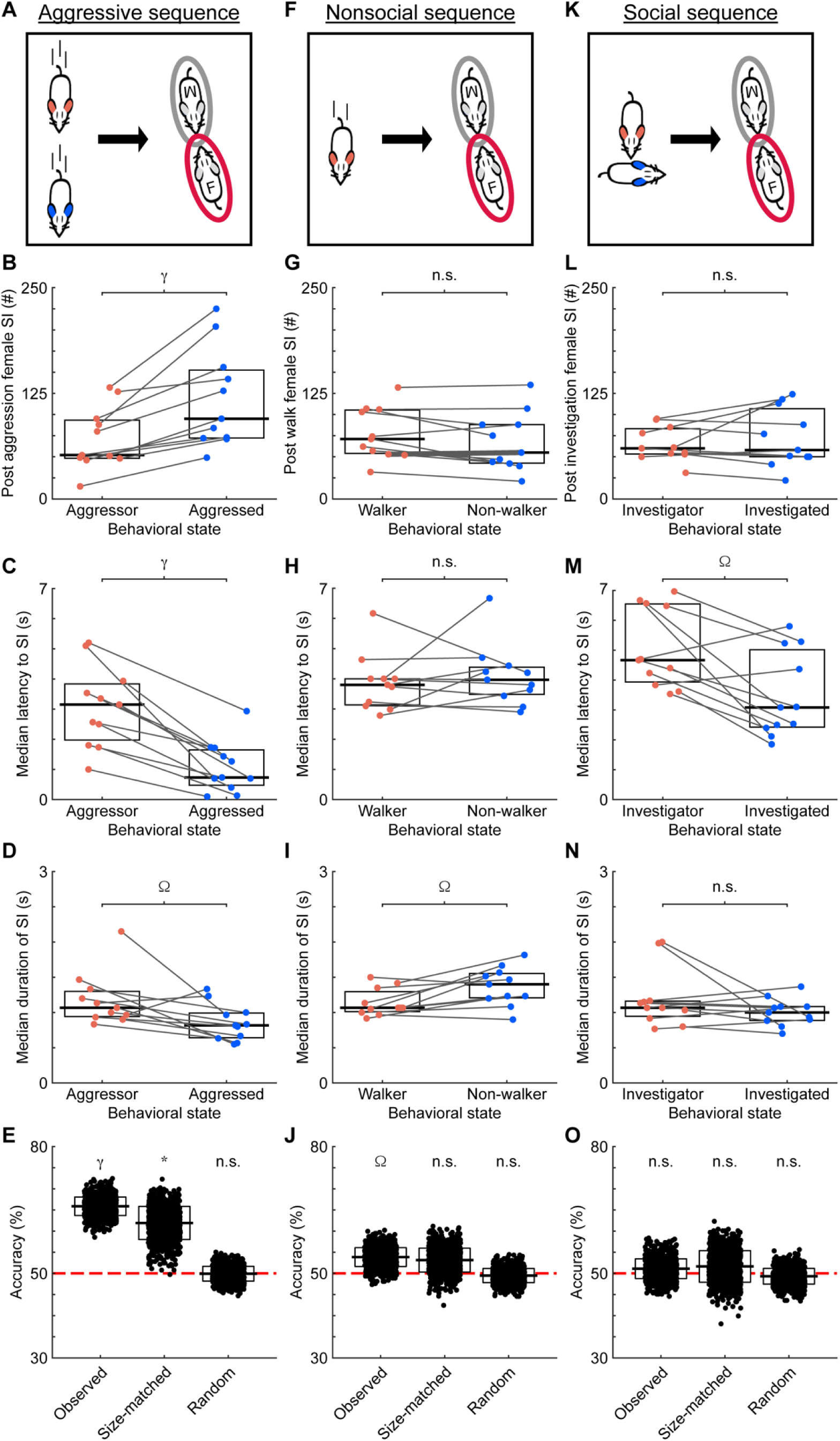
The behavioral state of an individual modulates subsequent interactions with females. (A) Schematic of aggressive social sequences. Sequences consisted of aggressive male interactions followed by male-female social interactions. (B) The number of male-female interactions following aggressive behaviors. Lines connect co-recorded mice. Black lines and white boxes show the medians and interquartile ranges (25-75%). (C) The latency between aggressive encounters and social interactions. (D) The duration of social interactions following aggressive encounters. (E) Performance of decoders when predicting the behavioral state of the male social partner in post-aggression social interactions. Black lines and white boxes show the means and standard deviations. The red line denotes chance levels. (F) Schematic of nonsocial sequences. Sequences consisted of a male walking in isolation followed by male-female social interactions. (G) As in B, for walking-triggered sequences. (H) As in C, for walking-triggered sequences. (I) As in D, for walking-triggered sequences. (J) As in E, for walking-triggered sequences. (K) Schematic of non-aggressive social sequences. Sequences consisted of investigative male interactions followed by male-female social interactions. (L) As in B, for investigation-triggered sequences. (M) As in C, for investigation-triggered sequences. (N) As in D, for investigation-triggered sequences. (O) As in E, for investigation-triggered sequences. Ω = p < 0.05, * = p < 0.01, γ = p < 0.001, n.s. = p ≥ 0.05

We used a machine-learning approach to further investigate the relationship between aggressive encounters and subsequent social interactions. Decision tree classifiers, which are predictive models, could decode the behavioral state of the male social partner (**Fig 3E**). Accuracies of the classifiers, trained on the latency to the social interaction and duration of the social interaction, ranged from 58.6% to 72.4% and significantly exceeded chance levels (chance = 50%; observed: 1-sided z-test, n = 1,000 iterations, z = 7.25, p < 0.001). When sample-size-matching the number of interactions the aggressor and aggressed had with females, decision tree classifiers could also predict the behavioral state of the social partner (1-sided z-test, n = 1,000 iterations, z = 3.04, p = 0.001). When randomizing the times of the hostile interactions between males that preceded the social interactions (**Methods**), the accuracies of the models dropped to chance levels (1-sided z-test, n = 1,000 iterations, z = - 0.05, p = 0.48).

We next asked whether an aggressive encounter between males triggered this phenomenon. To answer this question, we analyzed the temporal relationship between a non-social behavior, walking, and male-female social interactions (**Fig 3F**, **Methods**). When solitary walks preceded mixed-sex interactions, both the walker and non-walker were equally likely to interact with females (**Fig 3G**; walking male: median = 61, IQR = 67; non-walker: median = 59, IQR = 40; Wilcoxon Signed Rank Test, W = 47, p = 0.24). No differences between behavioral states were observed in the latency to interact (**Fig 3H**; walking male: median latency = 3.80 seconds, IQR = 0.73 seconds; non-walker: median latency = 3.90, IQR = 0.69 seconds; Wilcoxon Signed Rank Test, W = 21, p = 0.54). The duration of the interactions was significantly shorter for males that were not walking than walking males (**Fig 3I**; walking male: median duration = 1.33 seconds, IQR = 0.40 seconds; non-walker: median duration = 1.07 seconds, IQR = 0.38 seconds; Wilcoxon Signed Rank Test, W = 32, p = 0.68). Overall, the relationship between aggressive encounters and interactions differed from that of walks and interactions, suggesting that aggressive behavioral state mediates subsequent social interactions. Predicative models could successfully differentiate the behavioral state of the male social partner in non-social contexts (**Fig 3J**; 1-sided z-test, n = 1,000 iterations, z = 1.73, p = 0.04), but the accuracies of the predicative models dropped to chance levels when controlling for sample size and randomizing the times of the behaviors triggering the sequence (all 1-sided z-test, n = 1,000 iterations; size-matched: z = 1.11 p = 0.13; random: z = -0.31, p = 0.38).

Another possibility is that any male-male social encounter, not solely aggressive encounters, acts as a trigger for subsequent social interactions with females. Sequences in which non-aggressive social interactions (male investigations of males) were followed by opposite-sex interactions (**Fig 3K, Methods**) produced results mirroring walking-triggered sequences. Males were classified as investigating or being investigated based on which animal initiated the investigation. There were no differences between behavioral states when comparing the number of opposite-sex interactions (**Fig 3L**; investigating male: median = 60, IQR = 53.75; investigated male: median = 65, IQR = 26.25; Wilcoxon Signed Rank Test, W = 39, p = 0.62). While the latency between the investigation and the social interaction was shorter when the investigated male was the social partner (**Fig 3M**; investigating male: median = 4.67 seconds, IQR = 2.62 seconds; investigated male: median = 3.083 seconds, IQR = 2.59 seconds; Wilcoxon Signed Rank Test, W = 63, p = 0.001), the median latency for investigated males was higher than that of aggressed males. Furthermore, the duration of the male-female interactions following investigation was indistinguishable by behavioral state (**Fig 3N**; investigating male: median = 1.22 seconds, IQR = 0.58 seconds; investigated male: median = 0.93 seconds IQR = 0.28 seconds; Wilcoxon Signed Rank Test, W = 53.50, p = 0.07). Predicative model accuracies were at chance levels when attempting to identify the behavioral state of the male social partner in non-aggressive contexts (**Fig 3O**, all 1-sided z-tests, n = 1,000 iterations; observed: z = 0.47, p = 0.39; size-matched: z = 0.44, p = 0.33; random: z = - 0.37, p = 0.36). These results strongly suggest that an act of aggression triggers the aggressed male to interact with a female.

While the tendency to interact with a female following aggressive encounters appears to depend on an animal’s behavioral state, the hierarchical rank of an animal may also contribute to the observed pattern. To assess this possibility, we investigated the relationship between aggressive behaviors and male-female social interactions but accounted for aggregated aggression levels rather than the behavioral state. We found that the propensity of dominant and subordinate mice to interact with a female following a hostile interaction were similar (**S3 Fig A-B; S1 Table;** dominant median = 64, IQR = 80.25; subordinate median = 82, IQR = 71.25; Wilcoxon Signed Rank Test, W = 17, p = 0.17). There were no differences in latency between aggressive behavior and male-female interactions (**S3 Fig C; S1 Table;** dominant median = 1.91 seconds, IQR = 1.47; subordinate median = 1.33, IQR = 1.51; Wilcoxon Signed Rank Test, W = 41.5, p = 0.48) or the duration of social interactions (**S3 Fig D; S1 Table;** dominant median = 1 second, IQR = 0.55; subordinate median = 0.83, IQR = 0.33; Wilcoxon Signed Rank Test, W = 44.5, p = 0.33) when comparing aggregate aggressive levels. As seen in **S3 Fig**, predicative model accuracies were at chance levels when attempting to identify the behavioral state of the male social partner (**S1 Table**). When we performed the same analyses with aggregated aggression levels (dominant vs. subordinate) and examined non-social or non-aggressive triggers, we again found no differences during non-social (**S3 Fig F-J; S1 Table**) or non-aggressive triggered sequences (**S3 Fig K-O; S1 Table**). The only exception occurred when we predicted aggression levels for sample-size-matched investigation-triggered sequences (**S3 Fig O; S1 Table**). These findings suggest that behavioral state underlies the sequential nature of social interactions following aggressive encounters, not identity or aggregate aggression level.

To ensure that sample size did not bias the finding that aggressed males were more likely to engage with females after a hostile interaction, we performed a permutation test (**Fig 4A**, **Methods**). Here, we randomly selected 50 sequences from each recording and calculated a difference index between subsequent aggressed and aggressor interactions with a female. Values below zero indicate that following aggressive encounters, the aggressed males interacted with a female more than the aggressor. For aggression-triggered sequences, 100% of the permutations were below zero (2-sided z-test, n = 1,000 permutations, z = 7.85 p <0.001), suggesting that after hostile interactions between males, the aggressed male consistently interacts with the female first, and the effects are not due to sample size. For non-social (**Fig 4B**) and non-aggression (**Fig 4C**) triggered sequences, the distribution of indices was not significantly skewed towards aggressor or aggressed (2-sided z-test, n = 1,000 permutations each condition; non-social: z = -1.28, p = 0.21; non-aggressive: z = 0.45, p = 0.65). These analyses suggest that neither the sample size nor a subset of examples underlies the results.

**Fig. 4.**
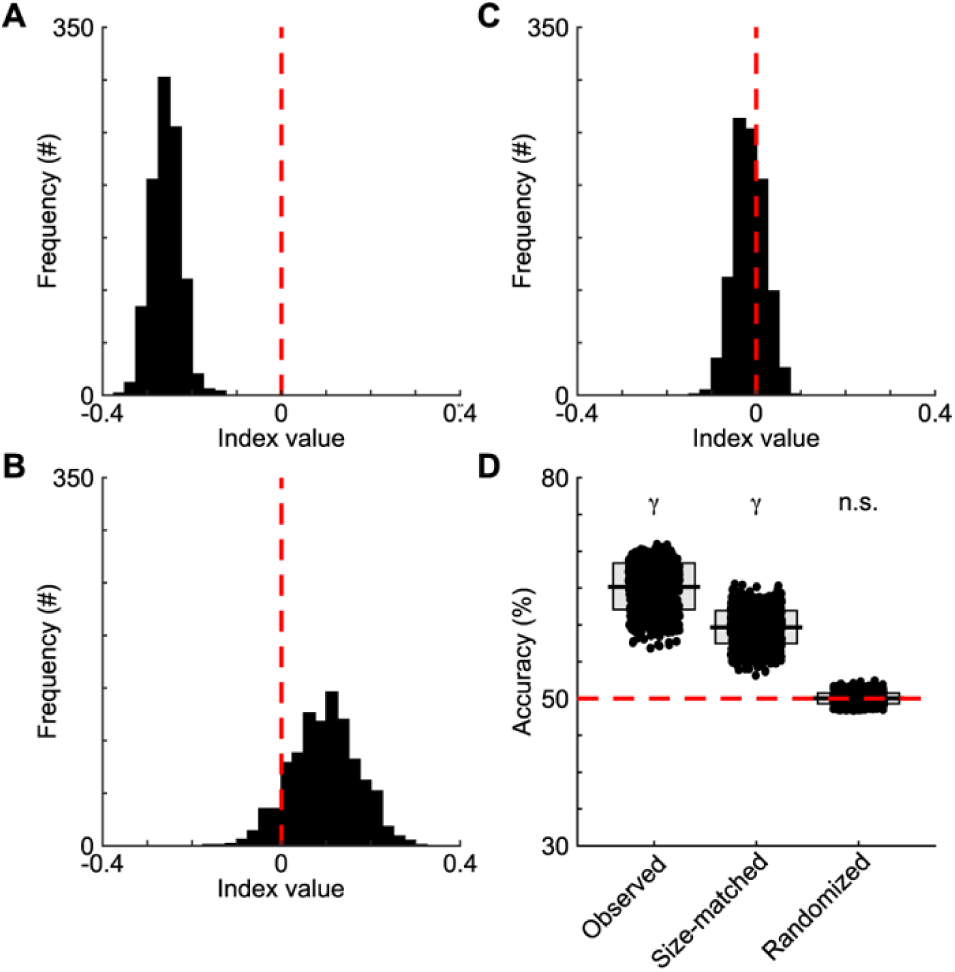
Aggressive-triggered sequences differ from walking- and investigation-triggered sequences. (A) A subsample of aggression-triggered sequences was randomly selected from each recording. Next, a difference index was calculated by subtracting the number of interactions between the aggressed male and female from the number of interactions between the aggressor male and female. The difference was then divided by the total to create an index. An index value below 0 reflects more interactions occurring between an aggressed male and female than an aggressor male and female. The random sampling procedure and index calculations were performed 1,000 times. Aggressed interactions with the female occurred significantly more often than aggressor interactions with the female, as the distribution was shifted to the left of zero. (B) As in A, for walking-triggered sequences, the indices were not significantly different from zero. (C) As in B, for investigation-triggered sequences. (D) Decoders’ performance when predicting aggressive- or nonaggressive-triggered sequences. The horizontal bars and boxes below the data show the means and standard deviations. The red line denotes chance levels. γ = p < 0.001, n.s. = p ≥ 0.05

Using decision tree classifiers trained on the latency between the preceding (aggressive, social, or non-social) behavior and the social interaction, the duration of the social interaction, and the behavioral state of the male social partner, we could predict the type of behavioral sequence (aggressive or control behavior) at rates higher than chance levels (**Fig 4D**; observed: 1-sided z-test, z = 4.73, p < 0.0001). The model’s accuracy did not depend on the number of examples, as we maintained high accuracy using equal numbers of aggressive and control sequences (**Fig 4D**; size-matched: z = 4.32, p < 0.0001). Additionally, accuracy was low when we attempted to decode sequence type on data where we randomized the timing of the aggressive, non-social, and non-aggressive behaviors while maintaining the times of the social interactions (**Fig 4D**; randomized: z = 0.05, p = 0.48). This evidence helps to substantiate the finding that a male’s behavioral state affects subsequent interactions with females after a hostile interaction.

The behavioral state of an animal plays a key role in the subsequent interactions with females. To address the importance of behavioral state, we preformed another permutation analysis. First, we randomized the identity of the males during aggressive encounters that preceded social interactions. Then we calculated a difference index between subsequent aggressed and aggressor interactions with the shuffled data. This procedure was performed 1,000 times to generate a distribution of index values. We then compared the actual index value (-0.25) to the shuffled distribution and found that it was significantly lower than the mean of the distribution (**Fig 5A**; mean of distribution = 0.0005, standard deviation = 0.02, z-score = -11.05, p < 0.0001). The subsequent interactions between the males and females were likely due to the males’ trajectories (**Fig 5B-C**). The angles between the male’s heading direction and the vector pointing from the male to the female was significantly different between the aggressor and aggressed animals (**Fig 5D-E**; aggressive state: median angle = 75.94°, variance = 21.77°; aggressed state: median = 41.34°, variance = 17.26°; Watson’s U2, U2 = 2.12, p < 0.001), with the angles of the aggressed skewed towards the female. The differences persisted when looking at individual animals (**Fig 5F-G**; aggressive state: median angle = 78.07°, variance = 2.74°; aggressed state: median = 40.72°, variance = 0.61°; Watson’s U2, U2 = 0.34, p = 0.002). The differences in the angles between the male’s heading direction and the vector pointing from the male to the female were not due to the orientation of the females relative to the males (**Fig 5H-K**). We found that the differences in the orientation of the aggressor and aggressed animals from the female at the end of the aggressive triggering event had no effect on the likelihood of interacting with a female, as there were no differences between aggressor and aggressed animals (**Fig 5L-M**; aggressive state: median angle difference = 79.59°, variance = 21.17°; aggressed state: median = 81.25°, variance = 20.68°; Watson’s U2, U2 = 0.05, p = 0.81). For individual animals, the differences in orientations between animals in an aggressed and aggressor behavioral states were similar (**Fig 5N-O**; aggressive state: median angle difference = 81.89°, variance = 0.68°; aggressed state: median = 82.31°, variance = 0.28°; Watson’s U2, U2 = 0.06, p = 0.57). These results suggest that heading direction of the aggressed male plays a role in the subsequent interactions with female, but the interactions were brief and frequent.

**Fig. 5.**
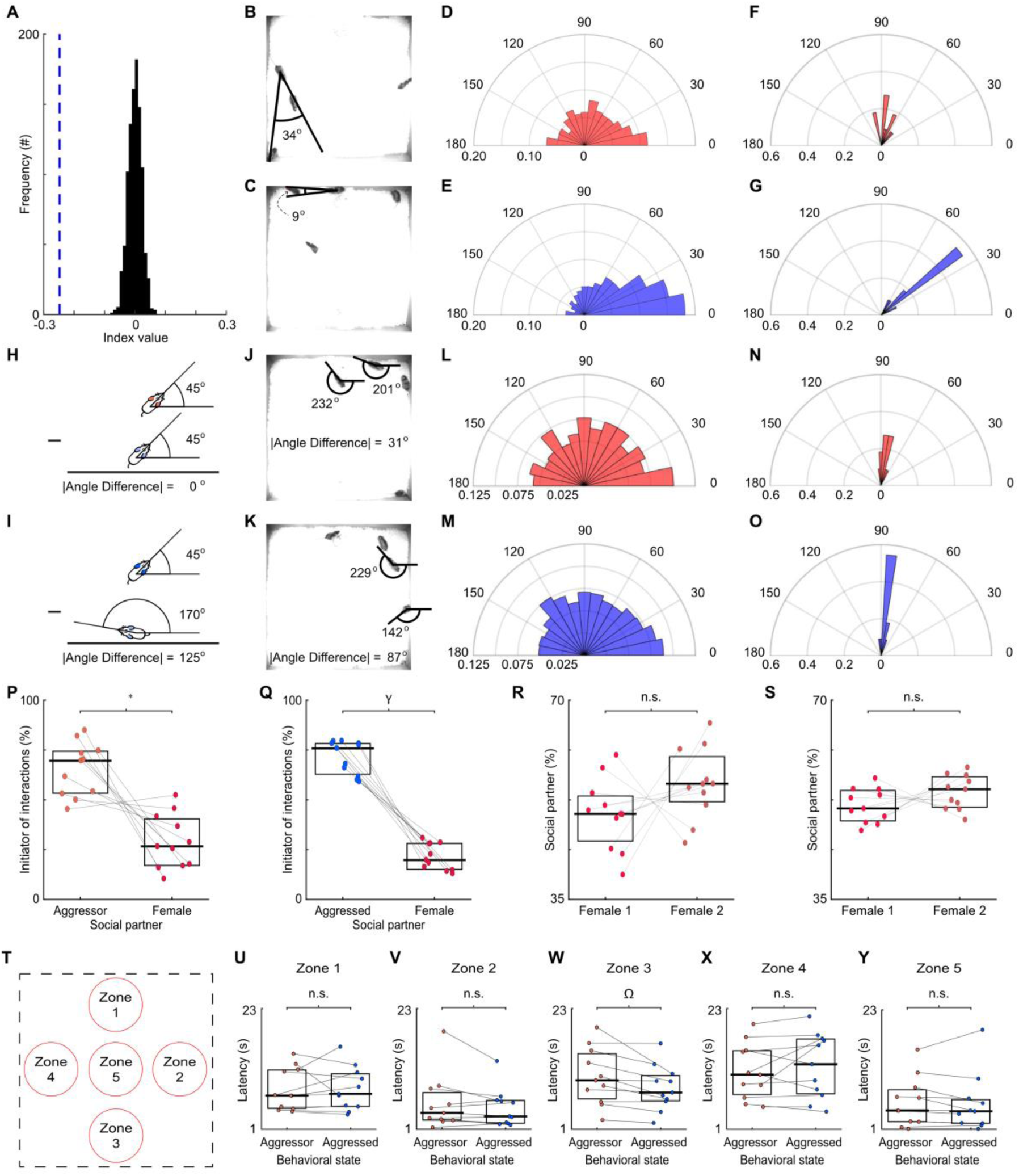
Features of aggression-triggered social interactions are dependent upon state, but not orientation of mice or spatial location in cage. (A) The identity of the aggressor was randomized, and we then quantified the propensity of animals to engage in an aggression-triggered social interaction. An index value was calculated with the shuffled data, as in Fig 4. This procedure was repeated 1,000 times to generate a distribution of index values. We compared the actual index value (denoted as a blue line) to the random distributions to generate a z-score. (B) Example showing the angle between the aggressor male’s heading direction and the vector pointing from the male to the female at the end of an aggressive encounter. (C) As in B, for the aggressed male. (D) Distribution of angles between the aggressor male’s heading direction and the vector pointing from the male to the female at the end of the aggressive encounter. Theta and radians on the polar plot represent angles in degrees and normalized frequency. Normalization was determined by dividing the number of occurrences in each bin by the number of post-aggression interactions between the aggressor and female. (E) As in D, for the aggressed male. (F) Median angle between the aggressor’s heading direction and the vector pointing from the animal to the female for individuals. (G) As in F, for the aggressed male. (H) Schematic showing absolute angle difference calculations for aggressor and female social partner. (I) As in H, for the aggressed male. (J) Example showing the angle difference between the orientations of aggressor male and female social partner. (K) As in J, for the aggressed male. (L) Polar plot showing distribution of angle differences at the end of aggressive behaviors when aggressor was the social partner in the aggression-triggered social interaction. (M) As in L, for the aggressed male. (N) Median angle difference for aggressor and aggressed. The horizontal bars and boxes show the medians and the interquartile ranges (25-75%). Lines connect co-recorded mice. (O) As in N, for the aggressed male. (P) Male-female social interactions initiated by the aggressor or female. Lines connect co-recorded mice. The horizontal bars and boxes below the data show the medians and interquartile ranges (25-75%). (Q) As in P, for interactions with the aggressed. (R) Proportion of interactions the aggressor male had with either female. Lines connect co-recorded mice. The horizontal bars and boxes below the data show the medians and interquartile ranges (25-75%). (S) As in R, for interactions with the aggressed. (T) Schematic showing arbitrary zones in the cage. (U) Median latency of aggressor and aggressed to reach zone 1 from the end of all aggressive behaviors. Lines connect co-recorded mice. The horizontal bars and boxes the data show the medians and interquartile ranges (25-75%). Significance tested using a Wilcoxon signed rank test. (V) As in U, for zone 2. (W) As in U, for zone 3. (X) As in U, for zone 4. (Y) As in U, for zone 5. Ω = p < 0.05, * = p < 0.01, γ = p < 0.001, n.s. = p ≥ 0.05

The male-female interactions following an aggressive encounter could be initiated by either females or males. To determine the initiator, we measured the instantaneous speeds and the positions of the social partners. Animals that were moving and getting closer to the social partner were considered the initiator. Regardless of behavioral state, opposite-sex social interactions following aggressive encounters were consistently initiated by males (**Fig 5P-Q**; median aggressor male proportion = 0.77, IQR = 0.26, Wilcoxon Signed Rank Test, W = 64, p = 0.003; median aggressed proportion = 0.79, IQR = 0.20, Wilcoxon Signed Rank Test, W = 69, p < 0.001), suggesting that onset of the social encounter following the aggressive behavior is driven by the aggressed males rather than by female social partners. Additionally, this phenomenon was not dependent on a particular female, as males in both behavioral states engaged in post-aggression social interactions with each female (**Fig 5R-S**; aggressor: female 1 median proportion = 47%, IQR = 9%, female 2 median proportion = 53%, IQR = 9%, Wilcoxon Signed Rank Test, W = 16, p = 0.15; aggressed: female 1 median proportion = 48%, IQR = 6%, female 2 proportion = 52%, IQR = 6%, Wilcoxon Signed Rank Test, W = 20, p = 0.08). The results suggest that males were initiating the majority of social interactions.

A possibility exists that the aggressed animal is escaping from the aggressor and encounters the female by happenstance. If this is the case, then the aggressed male should encounter any zones in the behavioral arena before the aggressor male. To evaluate this prospect, we quantified the latency to reach multiple arbitrary zones in the behavioral arena after an aggressive behavior ended. The zones were in the North central (zone 1), South central (zone 2), West central (zone 3), center (zone 4), and East central (zone 5) areas of the cage (**Fig 5T**). When analyzing each ach zone independently, we found no differences in latency to reach zones 1, 2, 4, and 5 (**Fig 5U-Y**; **Table 1**). These results add credence to our hypothesis that this phenomenon is state-dependent.

**Table 1.**
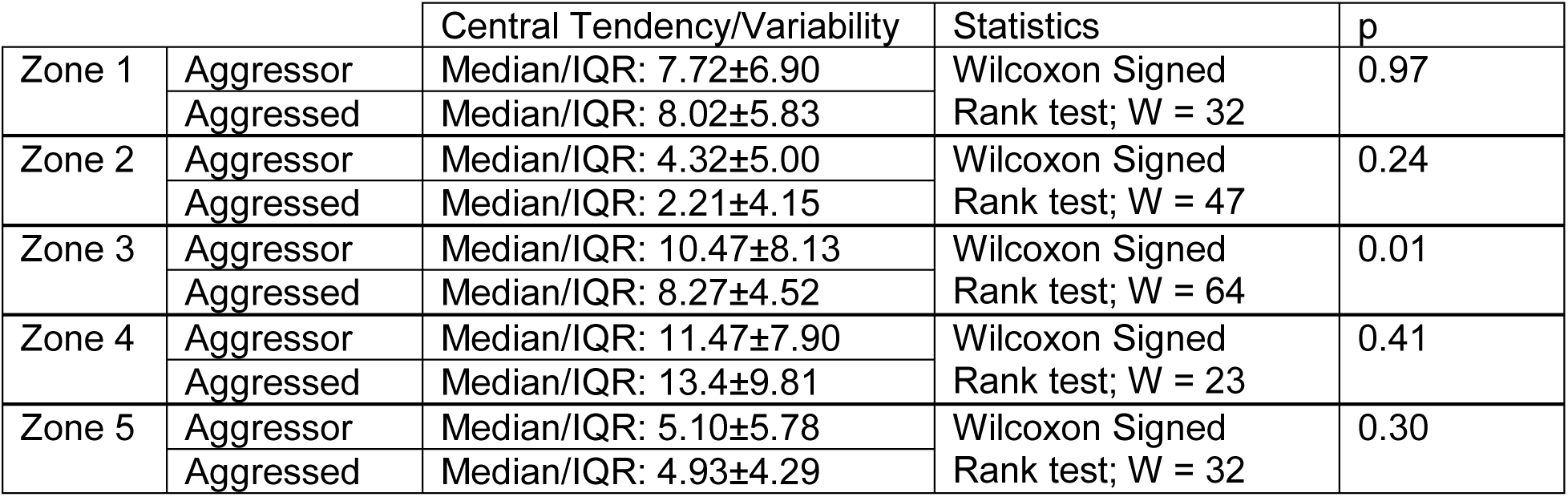
Statistical values corresponding to analyses presented in Fig 5U-Y.

### Temporal Profile of Aggression-Triggered Male-Female Interactions

Aggression-triggered social interaction sequences emerged early and persisted over time. The first aggression-triggered sequence in every recording occurred within 10.26 minutes (n = 11, median = 5.82 minutes, IQR = 3.78 minutes), whereas the last event occurred after 283.24 minutes (median = 297.27 minutes, IQR = 170.20 minutes). Over 5 hours, the proportion of sequences in which the aggressed animal interacted with a female was consistently higher than the aggressor (**Fig 6A**, **Table 2**). Using multi-class support vector machines (mcSVM) trained on the latency to the social interaction, duration of the interaction, and behavioral state, we assessed the accuracy in which the predictive models could identify the hour in which the sequence occurred. The accuracies of the models were below chance levels (**Fig 6B**, **Table 2**), suggesting that sequences were similar over time. The behavioral state could be predicted with decision tree classifiers for each hour of the recording (**Fig 6C**, **Table 2**, observed). However, when controlling for the number of sequences per hour, the accuracies of the models were significantly above chance levels only during the first hour (**Fig 6C**, **Table 2**, size-matched). When randomizing the hour in which the sequence occurred, the predictive model’s accuracy was indistinguishable from chance levels (**Fig 6C**, **Table 2**, randomized). These findings suggest that aggression-triggered female social interaction sequences occur at the onset and throughout the recordings.

**Fig. 6.**
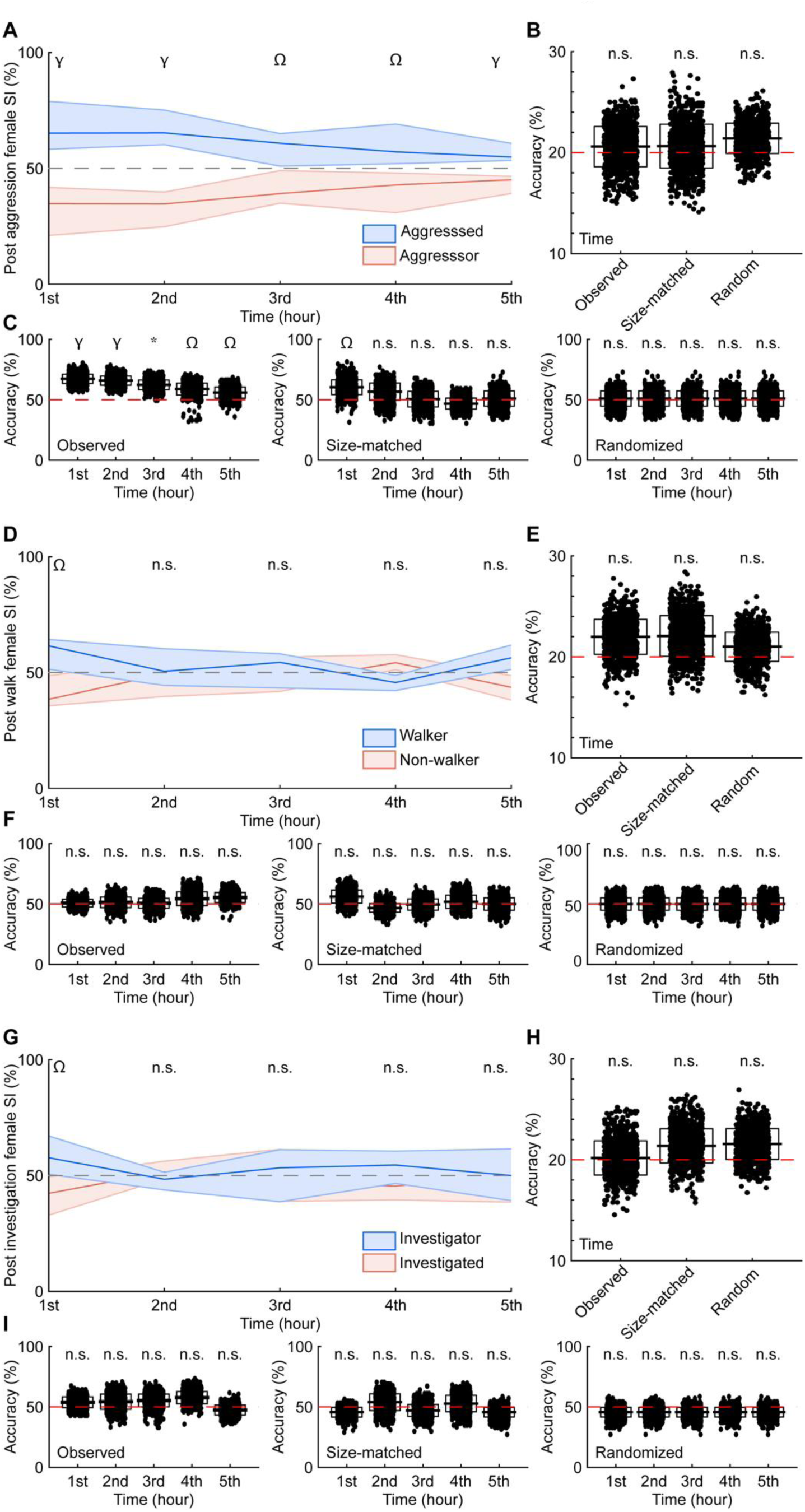
Aggression-triggered female social interaction sequences emerge early and persist over time. (A) The percentage of aggression-triggered sequences binned hourly for individuals in either an aggressed or aggressive behavioral state. The lines and shaded regions show the medians and interquartile ranges (25-75%). (B) Decoders’ performance when predicting when the sequences occurred. The horizontal bars and boxes below the data show the means and standard deviations. The red line denotes chance levels. (C) Decoders’ performance when predicting the behavioral state of the male interacting with the female for all observed data, size-matched controls, and randomized start times of aggressive male-male interactions. (D) As in A, for walking-triggered sequences. (E) As in B, for walking-triggered sequences. (F) As in C, for walking-triggered sequences. (G) As in A, for investigation-triggered sequences. (H) As in B, for investigation-triggered sequences. (I) As in C, for investigation-triggered sequences. Ω = p < 0.05, * = p < 0.01, γ = p < 0.001, n.s. = p ≥ 0.05

**Table 2.**
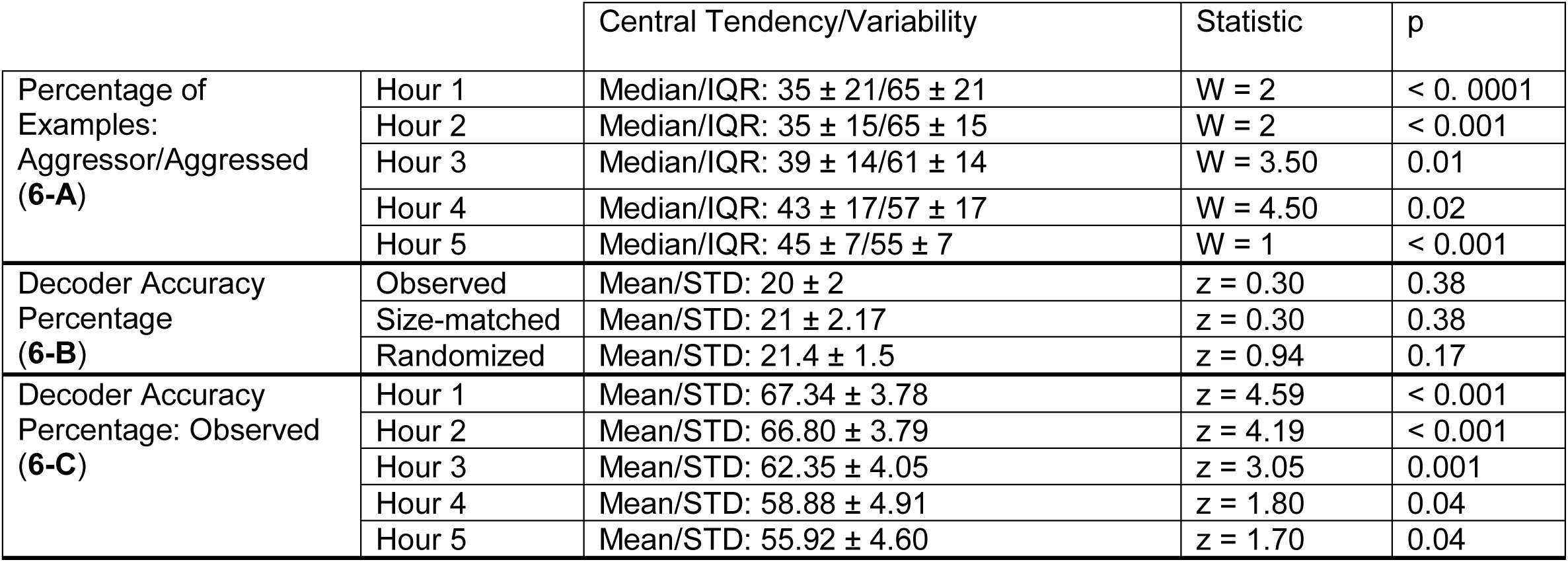
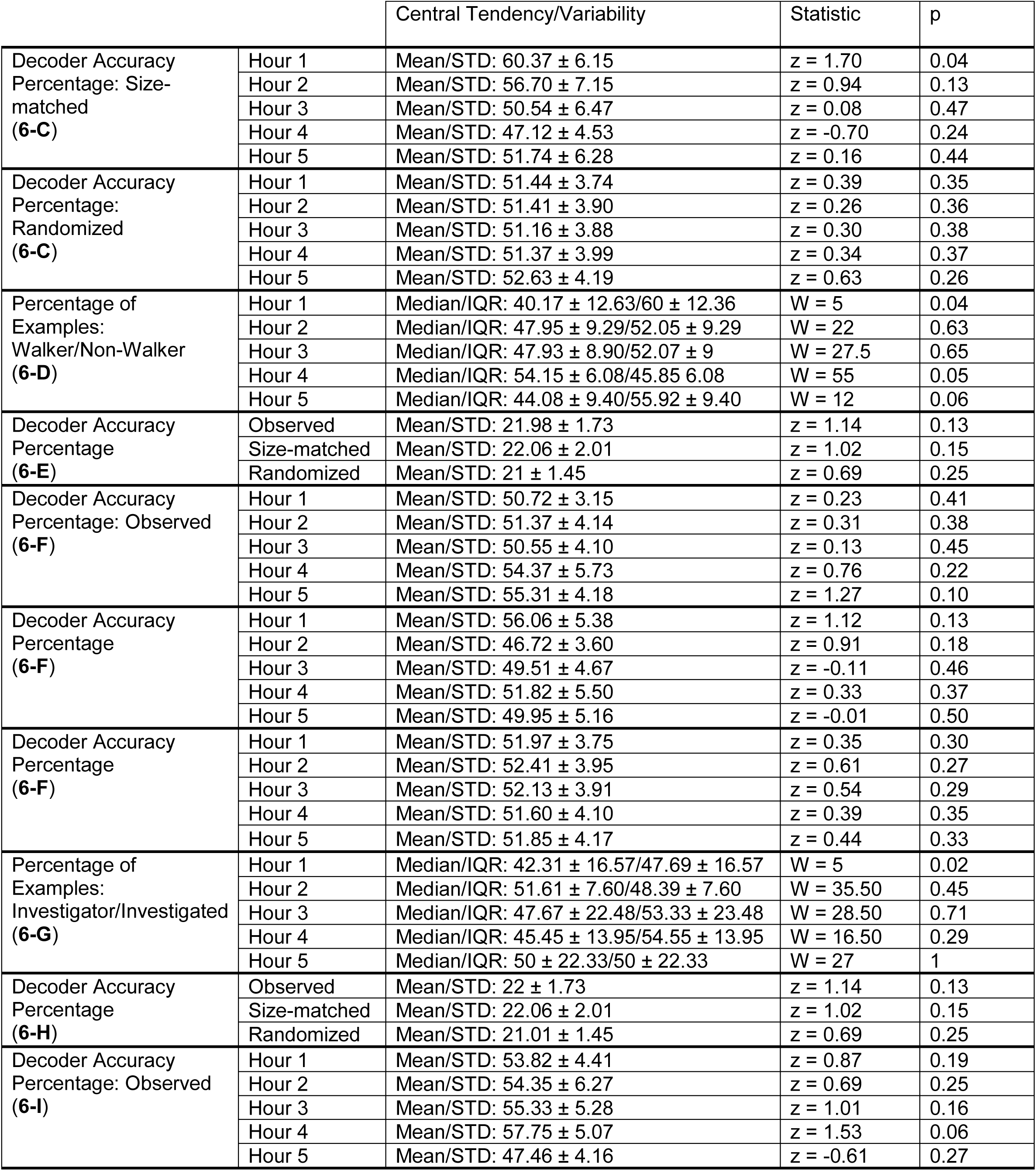
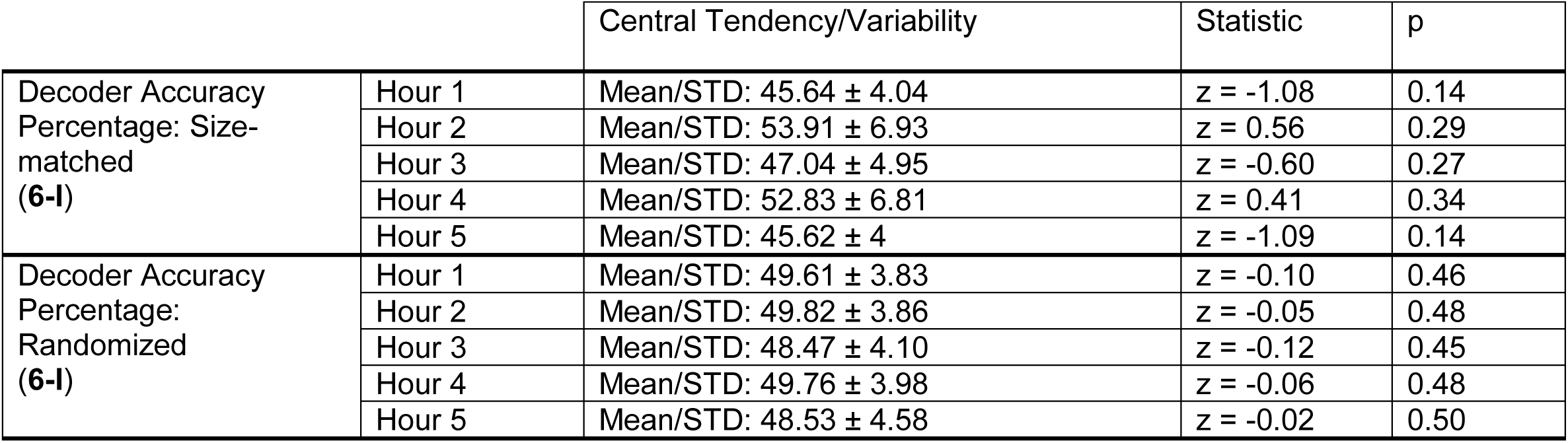
Statistical values for analyses presented in Fig 6. Bold values indicate corresponding figure panel.

The temporal dynamics of walking- and investigation-triggered female social interaction sequences were strikingly different from those of aggression-triggered sequences. The proportion of sequences in which the walking or investigating animal interacted with a female was consistently higher than the non-walking or investigated animal only in the first hour (**Fig 6D, G, Table 2**). After the first hour, no differences were observed in the walking- or investigation-triggered sequences. Our mcSVM models could not accurately decode when walking- or investigation-triggered sequences occurred (**Fig 6E, H, Table 2**). Additionally, multi-class support vector machines failed to decode walker/non-walker or investigator/investigated, regardless of the time (**Fig 6F, I, Table 2**), unlike decoders applied to aggression-triggered sequences (**Fig 6C**, **Table 2**). When accounting for aggregated aggression levels (dominant vs. subordinate), the proportion of sequences in which the dominant and subordinate animals interacted with a female was almost indistinguishable over time (**S4 Fig A, S2 Table**). The only observed differences occurred during the fourth hour of recording. Relative to chance, mcSVMs could not predict which hour the sequence occurred (**S4 Fig B, S2 Table**), and decision tree classifiers decoding the identity of the social partner had high accuracy only in the fourth hour when using all observed examples (**S4 Fig C, S2 Table**). These analyses further suggest that behavioral state specifically, rather than aggregate aggression level, is an important mediator of future social interactions throughout the behavioral experiments.

### Escape Mechanism for Deescalating Hostile Confrontations

Animals use multiple strategies to avoid confrontations, as injury and insult reduce biological fitness (6, 23). Thus, we reasoned that aggressed animals might purposefully interact with a female to divert the attention of the pursuing aggressor. To assess whether this was happening, we characterized the social interactions that followed the interactions between the aggressed male and a female. If this was a purposeful attempt to divert attention, one could reason that the male aggressor would engage with the same female. Indeed, this was the case (**Video 1**), as the majority of subsequent male and female social interactions occurred between the aggressive male and the female that previously engaged with the aggressed male (**Fig 7A**, Sequence type 1, 53%). Other types of social interactions also occurred after the aggressed male interacted with a female (**Table 3**). Sequence type 2, which involved the aggressed male interacting with the same female, occurred 21% of the time. Sequence type 3, social interactions between the aggressor and the other female, and type 4, social interactions between the aggressed and the other female, occurred 9% and 17% of the time, respectively. Sequence type 1 happened profoundly more often than the other sequence types (**Fig 7B**, Kruskal-Wallis with Dunn-Sidak post hoc correction; H(27.35), p < 0.001). Furthermore, sequence type 1 happened more often than expected by chance, as evidenced by bootstrap analyses (**Fig 7C**, **Table 4**). When training a mcSVM using the speed and distance traveled of the aggressor and aggressed animals during the aggressive behavior, we found that we could not distinguish the sequence types (**Methods**; data not shown, mean accuracy = 27%, standard deviation = 3%, 1-sided z-test, n = 1,000 iterations, z = 0.79, p = 0.43), suggesting that the aggressive states triggering the sequences are similar. Together, these findings show that aggressed males engage in distinct, reoccurring patterns of behavior after aggressive interactions, suggesting that they implement a “bait-and-switch” like tactic to escape hostile interactions.

**Fig. 7.**
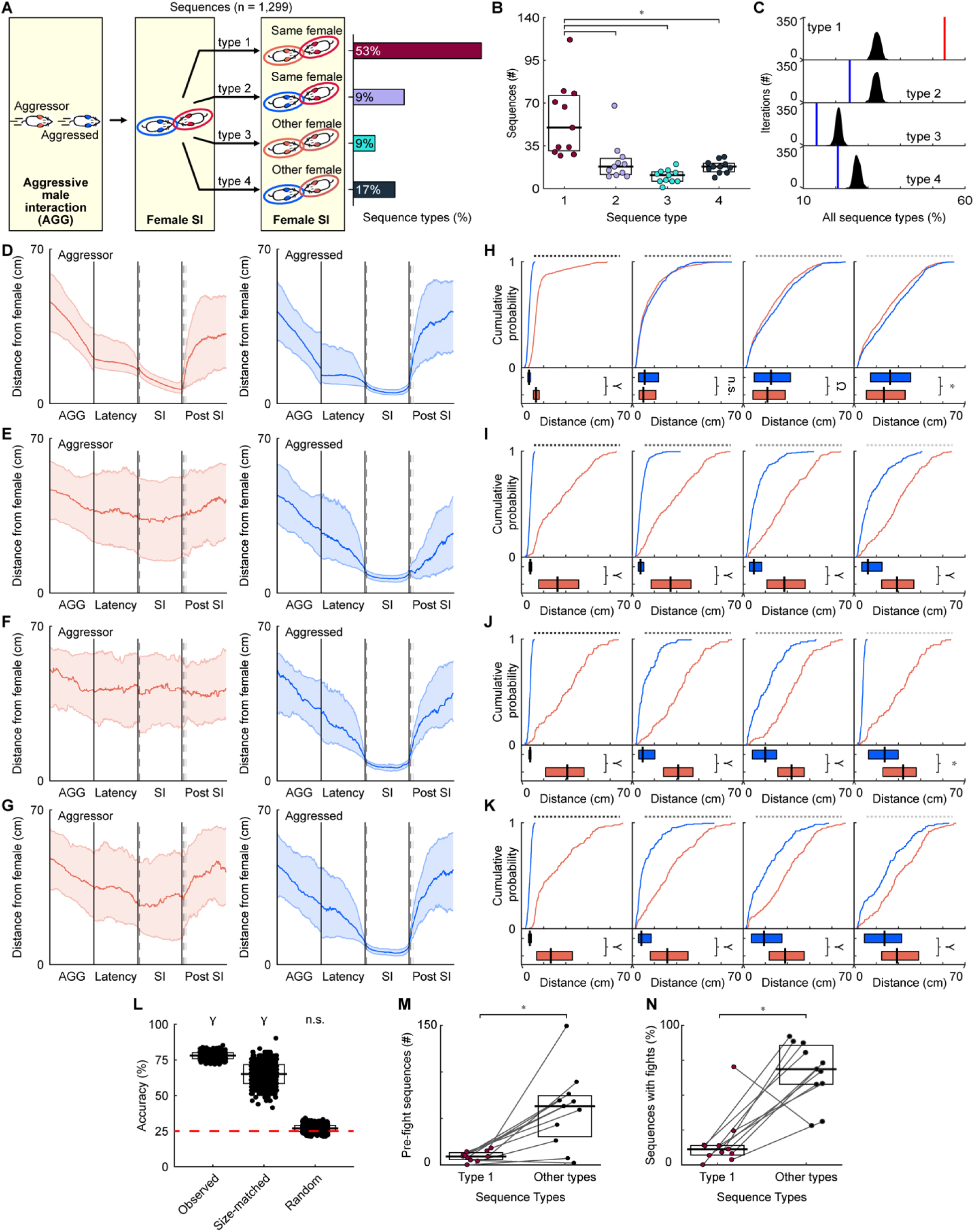
Submissive state-dependent behavioral strategies after aggressive encounters. (A) Schematic of sequence types and frequency of occurrence. (B) The number of each sequence type across recordings. The horizontal bars and boxes below the data show the medians and interquartile ranges (25-75%). (C) Difference between the percentage of sequence types and randomized distributions of sequences. Red and blue vertical lines denote percentages significantly above or below chance. (D-G) For every instance of each sequence type, distances between the aggressive or aggressed males and the female social partner were calculated during aggressive behaviors (AGG), the time between AGG and male-female social interactions, interactions, and 5 seconds post interaction. The lines and shaded regions show the medians and interquartile ranges (25-75%). The dashed lines indicate times representing one second after the start of the social interaction and post-interaction times of 1, 2, and 3 seconds. (D) Sequence type 1. (E) Sequence type 2. (F) Sequence type 3. (G) Sequence type 4. (H-K) Times for quantifying distances one second after the start of the social interaction and post-interaction times of 1, 2, and 3 seconds. Times correspond to the differently colored dashed lines in D, E, F, and G. Top: all distances. Bottom: median distances for each mouse. (H) Sequence type 1. (I) Sequence type 2. (J) Sequence type 3. (K) Sequence type 4. (L) Decoders’ performance predicting sequence type. The horizontal bars and boxes below the data show the means and standard deviations. The red line denotes chance levels. (M) Likelihood of fights occurring in the presence or absence of type 1 sequences. Lines connect co-recorded mice. (N) Percentage of sequences with fights. Lines connect co-recorded mice. Ω = p < 0.05, * = p < 0.01, γ = p < 0.001, n.s. = p ≥ 0.05

**Table 3.**
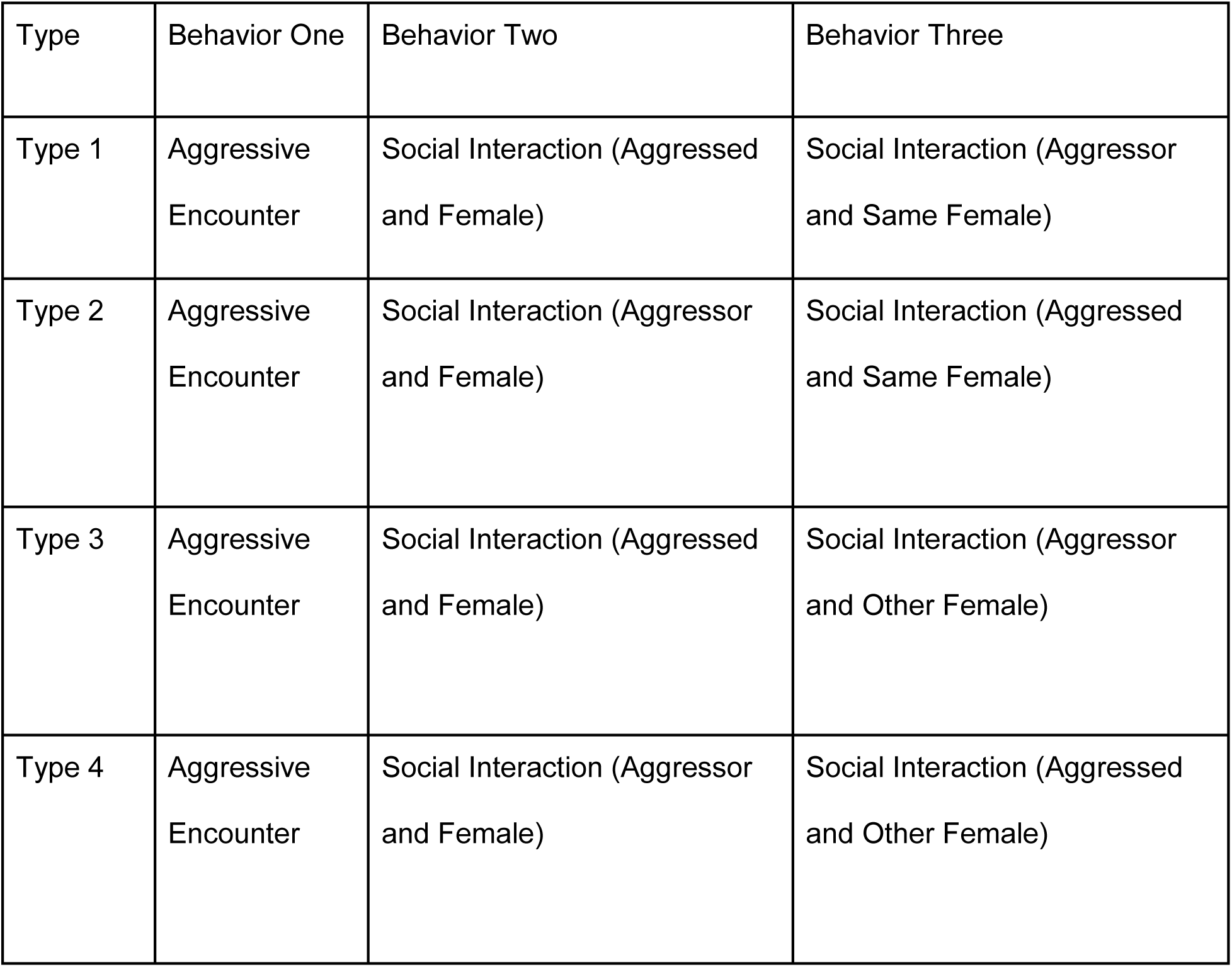
Description of sequence types.

| Type | Behavior One | Behavior Two | Behavior Three |
| --- | --- | --- | --- |
| Type 1 | Aggressive Encounter | Social Interaction (Aggressed and Female) | Social Interaction (Aggressor and Same Female) |
| Type 2 | Aggressive Encounter | Social Interaction (Aggressor and Female) | Social Interaction (Aggressed and Same Female) |
| Type 3 | Aggressive Encounter | Social Interaction (Aggressed and Female) | Social Interaction (Aggressor and Other Female) |
| Type 4 | Aggressive Encounter | Social Interaction (Aggressor and Female) | Social Interaction (Aggressed and Other Female) |

**Table 4.** Statistical values for analyses presented in Fig 7. Bold values indicate corresponding figure panel.

|  |  | Central Tendency/Variability | Statistic | p |
| --- | --- | --- | --- | --- |
| Difference from Random Distribution (7-C) | Type 1 | Mean/STD: 30 ± 1.15 | z = 20.05 | < 0.0001 |
|  | Type 2 | Mean/STD: 29.75 ± 1.21 | z = -7.44 | < 0.0001 |
|  | Type 3 | Mean/STD: 16.93 ± 0.98 | z = -7.58 | < 0.0001 |
|  | Type 4 | Mean/STD: 23.32 ± 1.12 | z = -5.86 | < 0.0001 |
| Distance Between Male and Female – Type 1: Aggressor/Aggressed (7-D, E) | Start + 1 second | Median/IQR: 13.13 ± 5.11/6.88 ± 2.52 | W = 192,814 | < 0.0001 |
|  | End + 1 second | Median/IQR: 9.99 ± 14.98/11.14 ± 18.04 | W = 71,772 | < 0.0001 |
|  | End + 2 seconds | Median/IQR: 21.82 ± 29.14/25 ± 32.75 | W = 73,311 | < 0.0001 |
|  | End + 3 seconds | Median/IQR: 26.78 ± 34.88/32.44 ± 36.30 | W = 79,583 | < 0.0001 |
| Distance Between Male and Female – Type 2: Aggressor/Aggressed (7-F, G) | Start + 1 second | Median/IQR: 32.61 ± 36.19/7.98 ± 2.45 | W = 29,521 | < 0.0001 |
|  | End + 1 second | Median/IQR: 34.44 ± 35.98/7.38 ± 5.03 | W = 29,028 | < 0.0001 |
|  | End + 2 seconds | Median/IQR: 37.07 ± 34.62/9.63 ± 11.05 | W = 28,036 | < 0.0001 |
|  | End + 3 seconds | Median/IQR: 38.77 ± 29.05/12.49 ± 18.25 | W = 26,449.5 | < 0.0001 |
| Distance Between Male and Female – Type 3: Aggressor/Aggressed (7-H, I) | Start + 1 second | Median/IQR: 41.30 ± 34.81/7.84 ± 1.85 | W = 6,210 | < 0.0001 |
|  | End + 1 second | Median/IQR: 41.43 ± 26.66/9.34 ± 14.61 | W = 5,980 | < 0.0001 |
|  | End + 2 seconds | Median/IQR: 43.60 ± 23.88/19.95 ± 21.85 | W = 5,702 | < 0.0001 |
|  | End + 3 seconds | Median/IQR: 44.32 ± 30.15/27.63 ± 26.92 | W = 4,995 | < 0.0001 |
| Distance Between Male and Female – Type 4: Aggressor/Aggressed (7-J, K) | Start + 1 second | Median/IQR: 26.53 ± 32.24/7.62 ± 2.60 | W = 19,306 | < 0.0001 |
|  | End + 1 second | Median/IQR: 31.58 ± 34.05/8.14 ± 11.39 | W = 16,730 | < 0.0001 |
|  | End + 2 seconds | Median/IQR: 37.76 ± 31.61/18.73 ± 27.50 | W = 5,358 | < 0.0001 |
|  | End + 3 seconds | Median/IQR: 38.71 ± 33.39/27.57 ± 33.722 | W = 14,307 | < 0.0001 |

**Video 1. Bait-and-switch sequence.**

Video shows a chase (aggressive behavior), a social interaction between the aggressed male and a female, and a subsequent interaction between the aggressor and the same female. The aggressor male’s fur is marked with 2 vertical stripes, while the aggressed is marked with 2 horizontal stripes. The female social partner is marked with a vertical slash. Video playback speed has been slowed to 15 frames per second.

If aggressed males were using the female as a means of diverting the attention of the aggressive male toward another animal and away from themselves, then we reasoned that the sequence of events would be reflected in the distance the males were from the female. Specifically, one would expect the aggressed male to be closer to the female during the initial social interaction and then move farther away as the aggressor approached. To quantify the behavior of the animals, we calculated the distance between the female in the first social interaction and both males (aggressor and aggressed) at multiple time points. The time points of interest were during the hostile interaction, the period between the hostile interaction and the first social interaction involving the aggressed mouse and the female, the first social interaction, and the next 5 seconds after the social interaction. **Fig 7D-G** shows the inter-individual distances between the male (aggressor or aggressed) and female social partner across all sequence types. For each of the sequence types, we observed notable differences between animals in the two behavioral states. We selected multiple time points to statistically compare the distances between males and females (**Fig 7H-K**, **Table 4**). The first time point was one second after the aggressed male started interacting with the female. For sequence type 1, the aggressed male was significantly closer to the female than the aggressor (**Fig 7H**, **Table 4**). One second after the social interaction ended, the distances between both males and the interacting female were equivalent (**Fig 7H**, **Table 4**). Two and three seconds after the social interaction, the aggressed male was significantly farther from the female than the aggressor (**Fig 7H**, **Table 4**). For sequence types 2-4, the aggressed male was always closer to the female than the aggressor (**Fig 7I-K**, **Table 4**).

We next wondered whether kinematic patterns predicted sequence type, as this could reflect that different sequences correspond with different behavioral strategies. We found that the distance between the female social partner and the males at six different time points (start of the aggressive encounter, end of the aggressive encounter, the start of the first social interaction, the end of the first social interaction, the start of the second social interaction, and the end of the second social interaction) could be used to decode sequence type using mcSVMs (**Fig 7L**; observed, 1-tailed z-test, n = 1,000 iterations, z = 25.03, p < 0.001). The relatively high number of type 1 sequences may have biased the classifier. As a control, we randomly selected the same number of each sequence type for each decoder iteration. When size-matching sequence types, classifiers still performed better than expected by chance (**Fig 7L**: size-matched, 1-tailed z-test, n = 1,000 iterations, z = 6.06, p < 0.001). Accuracy did not differ from chance when we randomized the sequence type label and used decoders to predict the sequence type (**Fig 7L**: randomized, 1-tailed z-test, n = 1,000 iterations, z = 0.93, p = 0.18). This evidence strongly supports the idea that the aggressed male might implement a “bait-and-switch” tactic to escape the hostile aggressor.

Escaping aggressive conspecifics and avoiding costly encounters is advantageous to an individual’s well-being (3). If aggressed animals successfully use a bait-and-switch like mechanism to escape hostile interactions and de-escalate social conflict, then fewer fights should occur between male interactions. Alternatively, the bait-and-switch could aggravate the aggressor and trigger an aggressive response, thus increasing the number of fights and escalating costly social conflicts. To address the two possibilities, we trained a supervised machine-learning classifier to detect fights (**Methods**). In total, 1,177 fights took place (n = 11 recordings, median = 88, IQR = 63.75) throughout the recordings (first fight, median time = 23.20 minutes, IQR = 24.69 minutes; last fight, median time = 296.62 minutes, IQR = 11.60 minutes). Compared to the other sequence types, fights rarely occurred when sequence type 1 (bait-and-switch) was present between aggressive male interactions (**Fig 7M**; n = 11, type 1: median = 9 fights, IQR = 7.50 fights, other types: median = 63 fights, IQR = 44 fights; Wilcoxon Signed Rank Test, W = 1, p = 0.002). Additionally, relative to the total number of fights, the proportion of fights following a bait-and-switch sequence was significantly lower than for the other sequence types (**Fig 7N**, n = 11, type 1: median = 10%, IQR = 6%, other types: median = 61%, IQR = 25%; Wilcoxon Signed Ranked Test, W = 3, p = 0.005). These findings suggest that the bait-and-switch sequence helps animals escape hostile interactions, and ultimately de-escalate conflict.

## Discussion

Animals consistently observe and monitor changes in their social environment, leading to adjustments in their behavior based on the integration of sensory feedback, previous social experiences, and internal states (3). Here, we implemented sophisticated, unbiased methods to quantify the behavior of males in groups of freely behaving mice and established aggressor-aggressed behavioral states as a centralized framework to examine natural social dynamics. We found that males use behavioral state-dependent strategies after hostile interactions to escape aggressive males and de-escalate confrontations. Specifically, we observed that after aggressive behaviors, the male in a submissive state was more likely to interact with a female social partner (**Fig 3**). Brief post-aggressive male-female social interactions occurred shortly after the aggressive encounter between males (**Fig 3**). This pattern appeared early, often, and throughout the five-hour behavioral recordings, highlighting the robustness of the phenomenon (**Fig 6**). Immediately after the interaction between the aggressed male and a female (first male-female interaction), we discovered that most of the subsequent male-female social interactions occurred with the female that the aggressed male interacted with and the aggressive male, suggesting that the aggressed males were implementing a bait-and-switch strategy (**Fig 7**). This potential behavioral strategy may serve to resolve agonistic encounters in a manner less costly to an animal’s fitness, as fights rarely occur after bait-and-switch sequences (**Fig 7**).

When examining sequences of behavior, we observed multiple aggression-triggered sequence types, with the bait-and-switch strategy (sequence type 1) occurring most frequently (**Fig 7**). The other sequence types were similar to sequence type 1 in that acts of aggression initiated the sequence, and an interaction between a male and a female followed the aggressive act. The four sequence types could be identified based on the second type of social interaction—the third step of the sequence. These interactions involved different possible combinations of social partners. Notably, the bait-and-switch strategy was rarely associated with fights (**Fig 7**), and we argued that this strategy was a mechanism to de-escalate conflict. However, the aggressive triggers initiating the sequences may reflect different modes of aggression. For example, animals in a more aggressive mode may focus on the target male. In contrast, animals in a less aggressive mode might be easily distracted and shift their interest to a female, leading to less fighting. Substantial evidence in both vertebrates and invertebrates indicates that behavioral states are differentiable by kinematics (24–28). Our predictive models could not discern sequence types based on the kinematics associated with the aggressive behaviors initiating the sequences. These findings suggest that the aggressive males initiating the sequence are in a similar behavioral mode. Nonetheless, future experiments that systematically measure behavioral states must be conducted to conclusively rule out the possibility that subtle differences in the aggressive triggers underlie the reduction in fights.

Animals escaping threats employ a diverse repertoire of defensive actions, including freezing, jumping, dashing, and avoiding the aggressor (29, 30). Defensive behaviors protect animals from danger and decrease the likelihood of future attacks (12). Threats, broadly speaking, also exist in predator and prey dynamics, and escape preludes survival. Although responses to social and predator pressures might manifest differently, individuals have adopted multiple strategies to protect themselves, and the approaches animals use to escape predation are well described (6, 9, 31–33). Some defensive actions involve reflex-like behaviors consistently initiated in response to predation. Crustacean tail flips are triggered in response to attacks from dragonfly nymphs and are typically the sole defense mechanism (34). Other animals, however, may need to optimize their defensive strategy depending on various environmental factors, including the availability of resources or the type of threat. To escape predation, schooling fish use uniform trajectories, while fish swimming alone vary their swimming angles (35). The direction frogs flee from predators depends on the position of the attack, as terrestrial attacks cause frogs to flee in the opposite direction, and aerial attacks cause frogs to move toward the predator’s flight path (36). If shelter is readily available, animals are more likely to flee to refuge than to freeze (7). While environmental cues are vital for optimizing defensive strategies against predation, animals must also assess trade-offs, as escaping may incur a loss of resources. For example, animals may flee their territory or give up mating opportunities to avoid predation (31). We observed that males in a submissive state sacrifice additional time with females, which theoretically could reduce their chances to copulate in favor of avoiding hostile interactions. We found that aggressed males—not the females— initiate social interactions following aggressive encounters but quickly move away from the female as the aggressor moves closer to her (**Fig 7**). This movement pattern suggests that aggressed males may use the bait-and-switch strategy to distract the aggressor with a potential mating opportunity. Distraction to avoid adverse consequences also occurs in humans; to avoid being targeted by a bully, individuals often engage in bullying behavior toward perceived lower-ranked classmates (37). The bait-and-switch tactic implemented by male mice in a submissive state could reflect the ability of animals to incorporate information about their surroundings and make optimal choices to avoid threats, as mirrored in other species. These findings suggest that mice integrate information about their behavioral state and that of others to shape dynamic social interactions and implement behavioral strategies.

Hierarchical rank or social status regulates animal interactions in multiple species, from wasps and fish to humans and other primates (38, 39). Switching between behavioral states— aggressor (the animal acting aggressively) or aggressed (the recipient of aggression)— decreases after establishing hierarchical rank (40). When agonistic interactions occur in an established hierarchy, dominant animals act as the aggressor to reinforce their social status (41). These agonistic encounters foreshadow a predictable sequence for dominant and submissive animals (42). Dominant animals initiate the sequence with a threat display and culminate with physical attacks (43, 44). Submissive animals must detect the threat, initiate a behavioral response, execute an escape behavior, and terminate the response when safe (6, 11). In some species, an animal’s social status may affect escape strategies (15, 45). For instance, changes in social status in crayfish alter the type of avoidance reaction displayed during aggressive encounters (46). Social rank also informs how individuals respond to chronic psychosocial stress. Larrieu et al. (47) showed that dominant mice develop an avoidant-like phenotype after chronically being defeated, suggesting prior experience impacts behavioral strategies. Our results indicate that the behavioral state of an animal rather than aggregate aggression levels influence escape strategies.

Our findings show that males engage in distinct behavioral patterns after aggressive encounters, and these sequences are associated with fewer fights. We interpreted this as a possible bait-and-switch strategy that males perform in a submissive state to de-escalate a confrontation. However, our experimental design prevented a thorough investigation of the underpinnings of this behavioral phenomenon as we sought to analyze naturalistic behavior. This study, instead, describes a frequently occurring behavior observed in males and paves the way for future experiments facilitating a mechanistic understanding of the potential escape strategy. Our studies also included small, sex-balanced groups, which may not represent groups of animals living outside of laboratory conditions. Future studies are required to manipulate the number of animals and combinations of males and females, which would allow a broader understanding of how practical the potential strategy is to conditions outside of laboratory settings. Our study also focused on animals without a clear social hierarchy. Because social status affects behavioral decisions (48–50), animals with established social ranks may engage in different behavioral sequences. Likely, the bait-and-switch sequence would not be observed in groups with an established pecking order, as hostility decreases with hierarchy formation (51), and aggression triggers these sequences, but this scenario should be tested further. Here, we characterized a behavioral sequence in which the female serves as a more potent distractor and a priority over the aggressed male. However, several questions remain. For instance, can males distract aggressors with objects or female scent cues? Females also form social hierarchies (52) and adopt strategies to escape unwanted attention from males (52–54). A distinct possibility exists that female mice also implement a similar approach for escaping aggressive interactions, and future experiments should fully characterize female behavior and determine if this phenomenon is also observed in females. Regardless of the limitations, the bait-and-switch sequence represents a discrete state-dependent behavioral strategy that requires animals to incorporate social information to guide behavioral decisions. Additionally, this strategy provides a framework for studying flexible, socially relevant behavioral strategies in laboratory settings and the underlying neural mechanisms.

## Methods

### Experimental Model and Subject Details

The data presented here were previously published (17), but all the analyses described were specific to the current study. The University of Delaware Animal Care and Use Committee, which complies with standards from the National Institutes of Health, approved all experimental protocols. Adult (13-21 weeks) male (n = 22) and female (n = 22) B6.CAST-Cdh23Ahl+/Kjn (stock number 002756, Jackson Laboratory, Bar Harbor, ME) mice were used. Mice were housed in a humidity-controlled, temperature-regulated colony room at the University of Delaware and were kept on a 12-hour light/dark cycle (lights off at 19:00 EST). At three weeks of age, mice were weaned and genotyped. Tail samples were sent to TransnetYX, and only mice expressing *Cdh23* were used in behavioral experiments. Mice were individually tagged with a light-activated microtransponder (p-Chip, PharmaSeq) subcutaneously implanted at the base of the tail. After weaning, animals were group-housed with same-sex siblings (3-5 mice per cage). All cages contained ALPHA-dri bedding and environmental enrichment. Mice had *ad libitum* access to food and water. Mice used in the experiments were age matched. Males were size matched, as were the females.

### Software and Algorithms

Matlab 2013, Matlab 2014, Matlab 2016

MOuse TRacker (MOTR, https://motr.janelia.org) (16)

Janelia Automatic Animal Behavior Annotator (JAABA, https://jaaba.sourceforge.net) (18)

### General Experimental Design

At least two weeks prior to the behavioral experiment, mice were isolate-housed to minimize the effects of group housing and hierarchal rank on social behavior (55, 56). For identification purposes, mice were painted with a unique back pattern using non-toxic hair dye (Clairol Nice ‘N Easy, Born Blonde Maxi) under light anesthesia at least two days before to the recording (16). Every mouse was randomly assigned and painted with a back pattern consisting of five dots, one diagonal slash, two vertical lines, or two horizontal lines. The day after painting, mice underwent a 10-minute exposure to a mouse of the opposite sex to increase social communication (57, 58). The opposite-sex partners were not used in behavioral recordings but were repeatedly used with multiple test subjects during the opposite-sex exposure sessions. If animals attempted to copulate during the opposite-sex exposure session, a trained observer ended the session. Two hours before the anticipated start of the recording, the estrous stage of the female mice was assessed using non-invasive vaginal lavage and cytological assessment. Specifically, cells were collected with a saline wash, placed on a slide, stained with crystal violet, and examined under a light microscope (VWR, Product number: 89404-890). Pictures of cells were taken using a camera (World Precision Instruments, Product number: USBCAM50) attached via a coupler (World Precision Instruments, Product number: 501381) to the microscope. Females were determined to be in estrus if their cells were predominantly cornified squamous epithelial cells lacking a nucleus (59). Recordings were only conducted if both females were in estrus. Otherwise, estrous testing continued until both females were in estrus.

For each recording, two males and two females were housed together for 5 hours in a mesh-walled cage (9218T25, McMaster-Carr), surrounded by Sonex foam (VLW-35, Pinta Acoustic). Mice were recorded for 5 hours to ensure adequate sampling of behaviors and to allow us to perform temporal analyses. The cage was placed in a custom-built anechoic chamber. For three of the eleven recordings, the cage was a cylinder (height – 91.4cm; diameter – 68.6), and for the remaining eight recordings, the cage was a cuboid with a frame (width, 76.2 cm; length 76.2 cm; height, 61.0 cm; **Fig 1B, C, S1 Fig**). The frame was made of extruded aluminum (8020). For the cylindrical cage, the mesh was secured between two plastic rings surrounding the arena, with one pair at the top of the cage and the other at the bottom of the cage.

Video data were continuously recorded (30 frames per second) for the entire five-hour experiment using a camera (GS3-U3-41C6M-C, FLIR) and BIAS software. Custom-written software controlled and synchronized the camera. The video data were stored on a PC (Z620, Hewlett-Packard). Infrared lights (IR-LT30, GANZ) were positioned above the cage and used to illuminate the arena to aid in tracking the mice. Before mice were placed in the cage, the cage was filled with ALPHA-dri bedding to an approximate depth of 0.5 inches. ALPHA-dri bedding was used to increase the color contrast between the cage floor and the mice. Each mouse was recorded individually for 10 minutes after the five-hour recording to aid with automated tracking (16).

### Data Processing

We used a data analysis pipeline set up on the University of Delaware’s FARBER computer cluster (http://docs.hpc.udel.edu) to determine the trajectory of each mouse. We used a program called MOTR to track the mice. Briefly, this program fits an ellipse around each mouse for every video frame. MOTR then determines the x and y position of the ellipse’s center, the orientation of the ellipse, and the size of the ellipse’s semi-major and semi-minor axes. This program also allows us to calculate nose position, distance from other animals, and instantaneous speed for each mouse at every frame of video. After the trajectories were determined, tracking was visually inspected by a trained observer.

### Automatic Extraction of Social Behaviors

We used JAABA, a machine-based learning program, to extract behaviors. Behaviors were manually classified based on definitions from previous work (17) (**Table 5**). Here, we focused on behaviors with clearly identifiable behavioral states. Aggressive social behaviors included chasing and fleeing, non-social behavior behaviors included walking, and non-aggressive social behaviors included male-male investigation. Behavioral states were identified as the following: 1) aggressor (the male chasing or the male being fled from) and aggressed (the male being chased or the male fleeing) for social aggressive, 2) walking or not walking for non-social, and 3) investigating and investigated for social non-aggressive. We also identified fights, but the aggressor was not discernable, therefore, these behaviors were not included in social aggressive behaviors, as we were specifically interested in assessing behavioral roles. All behaviors had a false positive rate below 5%, as determined by manual ground truthing (17).

**Table 5.** Behavioral definitions used to create JAABA classifiers.

| Behavior Name | Behavior Definition | Number of Examples | Post Hoc Refinements |
| --- | --- | --- | --- |
| Male Chase Male | A male follows another male while the two mice are within two body lengths of each other | 2,562 | Duration > 6 frames<br>Trajectories of males overlapping by at least 20%<br>Confidence scores > 0.5<br>Distance between males < 30 cm<br>Distance between chased animal and closed female > 15 cm |
| Male Being Chased | The male that is being followed | 2,562 | Same as Male Chase Male |
| Flee | A male running away from the other male | 851 | Duration > 10 frames<br>Confidence Score > 0.7<br>Closest animal < 8cm |
| Fled From | The male that the fleeing male is escaping; this male is typically stationary | 851 | Same as Flee |
| Walk | A mouse moves around the cage, in isolation (i.e., no other mouse within 35 cm) | 21,953 | Duration > 20 frames<br>Average Score > 1 |
| Male Investigate Male | Two mice touching, usually including sniffing. Can include nose to body, nose to nose, and/or anogenital investigation. This excludes other defined behaviors. | 4,661 | Confidence Score > 0.11<br>Speed < .19 & > .015<br>Average distance between males > 0.04 cm |
| Fight | Both males engaging in physical contact. Involves biting, wrestling, and rolling over each other. | 1,177 | Duration > 30 frames<br>Confidence Score > 1.5<br>Distance between males and nearest female at start > 5 cm<br>Average speed > 7.5 |

To assess differences in aggression across recordings, we calculated an aggression score (**Fig 2A**). The number of aggressive behaviors male one performed was subtracted from the number of aggressive behaviors male two performed. This value was divided by the total number of aggressive behaviors. Note that the labels of males 1 and 2 are arbitrary. Males with a higher number of behaviors were categorized as the dominant, and males with a lower number were categorized as the subordinate (40). Fights were not included in the aggression score because each male was in the same behavioral state (aggressor).

### Quantifying Social Interaction

A custom-written Matlab script was used to quantify social interaction. Pairs of mice were considered socially interacting when they were separated by less than one body length (≤ six centimeters). Thus, we used the mouse’s centroid position, major axis, minor axis, and heading direction (each value was calculated by MOTR) to automatically fit a “social ellipse” around each animal. Ellipses were fitted around each of the four animals for every video frame. The social ellipse extended three centimeters in front and behind the animal. Through manual investigation, we found that social interactions lasting fewer than six frames appeared to be interactions in which the mice quickly moved past each other. Therefore, all social interactions were required to last at least six frames. At every video frame, we determined whether a male was socially interacting with a female. Social interaction was defined as periods when ellipses were overlapping (**Fig 2E**).

We relied upon the animals’ instantaneous speeds and their physical positions relative to each other to automatically determine which mouse initiated each interaction. Using manual observation, we determined that when instantaneous speed was less than 0.023 cm/second, mice engaged in stationary motor behaviors, such as grooming and not moving towards or away from other animals. Mice with instantaneous speeds less than 0.023 cm/second at the start of social interactions were considered stationary. Through manual observation, we found that interactions were typically initiated by the other mouse when one mouse was stationary. Therefore, if one mouse was stationary in an interacting pair, the other mouse was classified as the initiator. If the instantaneous speed of both mice exceeded 0.023 cm/s, we determined which animal initiated the interaction by quantifying when the front of the ellipses began to overlap. We used the nose position to determine an animal’s heading direction and the front of the ellipse. We then assessed which ellipse pierced the other ellipse. However, if both animals were heading toward each other and the fronts of the ellipses overlapped within six frames, the initiation was classified as a mutual initiation. We confirmed the accuracy of the initiation software by comparing its results with those of a trained human observer. Results showing which animal initiated the social interaction are shown in **Fig 5P-S**.

### Sequences of Behavior

#### Aggressive Social Behaviors

To explore the dynamics between aggressive behaviors and male-female social interactions (**Fig 3A-E**), we found the start and end times of the aggressive behaviors and the male-female social interactions. We also identified the actors in aggressive behaviors, and social interactions. Behaviors were organized temporally, and we identified all sequences in which a male-female SI followed an aggressive behavior, termed aggression-triggered SIs. We first quantified the number of times males in either behavioral state were the male participants in the aggression-triggered SI. We statistically compared these values using a Wilcoxon Signed Rank test as the data did not meet the assumptions for normality (determined using the Kolmogorov-Smirnov test and visual inspection). We determined the latency between the aggressive behavior and the SI by subtracting the start time of the SI from the end time of the aggressive behavior. Examples were included even if the social interaction began before the end of the aggressive behavior. The duration of the SI was calculated by subtracting the SI’s end time from the start time. Within each recording, we found the median SI duration and latency between aggressive encounter and SI for both the aggressor and aggressed across all aggression-triggered SIs in which they participated. Median values from all recordings were compared using a Wilcoxon Signed Rank test. We also compared the number of sequences, median latency, and median duration using aggregate aggression levels rather than behavioral states to categorize the male social partner using Wilcoxon Signed Rank tests (**S3 Fig A-E**). To demonstrate that our results were state-dependent, we performed a permutation analysis in which we randomized the aggressor identity, while keeping the order and duration of all behaviors constant. We then determined the number of events where the randomized aggressor or aggressed engaged in the subsequent interaction. Then, we calculated a difference index using the following formula:

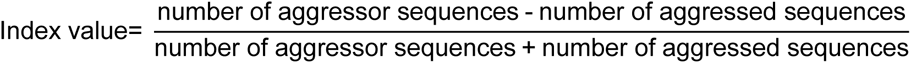

This procedure was repeated 1,000 times for each recording to generate a distribution of index values. After calculating the actual index value for the dataset, we computed a z-score using the following formula (**Fig 5A**):

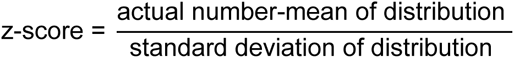

To compare the angles between the heading direction of the male and the vector pointing from the male to the female (**Fig 5B-G**), we used three coordinates (generated from the output of our tracking program, MOTR: the center of the male’s body, the nose of the male, and the center of the female’s body). The vector representing the male’s heading direction (v1) was determined using the center of the male’s body and nose. The vector between the male and female (v2) was calculated using the center of the male and female’s body. To determine the angle, we used the following formula:

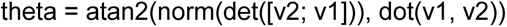

where the Matlab functions atan2, norm, det, and dot represent the four-quadrant inverse tangent, vector normalization, matrix determinant, and the dot product. To calculate the difference in the angle of ellipses (extracted by MOTR) between the social partners (aggressor/aggressed and female), we used the Matlab angdiff function (**Fig 5H-O**). To assess statistical differences in circular data, we used a Watson’s U2 test (<u>Matlab File Exchange</u>). The circular median and variance were calculated with the following functions downloaded from the <u>Matlab circular statistics toolbox</u>: circ_median and circ_var. Additionally, we compared the proportion of events where males (aggressor or aggressed) interacted with each female relative to the total number of sequences observed (**Fig 5P-S**). Again, comparisons were made using a Wilcoxon Signed Rank test.

To further demonstrate that our results were state-dependent, we quantified an animal’s latency to reach multiple arbitrarily defined zones after each aggressive behavior (**Fig 5T-Y**). In each recording, we computationally created circles with radii of 6 centimeters at 5 different locations. The locations included the North central (zone 1), South central (zone 2), West central (zone 3), center of the cage (zone 4), and East central (zone 5) areas of the cage. Each location was independently analyzed. We used tracking information to determine when the nose of each male crossed the boundary of the circle relative to the end time of each aggressive behavior. These values were then averaged across all events. We then used a Wilcoxon Signed Rank Test to compare aggressor median latency to aggressed median latency.

#### Non-Social Behaviors

To assess the relationship between non-social behaviors and social interactions, we repeated the aggression-triggered SI analyses, but used walking-triggered SIs (**Fig 3F-J**). Because the number of walks exceeded the number of aggressive behaviors in each recording, we randomly selected a subset of walks to match the number of aggressive behaviors. The male walking immediately before the SI was classified as a walker, while the male not walking was classified as a non-walker. We statistically compared the number, median latency, and median duration of sequences in which the walker participated to sequences in which the non-walker participated using a Wilcoxon Signed Rank test. Like aggression-triggered SIs, we also compared dominant and subordinate values by using aggregate aggression levels rather than behavioral states to categorize the male social partner (**S3 Fig F-J**).

#### Non-Aggressive Social Behaviors

To ensure that effects observed during aggression-triggered SIs were specific to aggressive behaviors and not social behavior more broadly, we implemented a social control in which we analyzed investigation-triggered SIs (**Fig 3K-O**). In all but one recording, there were more investigations than aggressive behaviors. For the recording with fewer investigations, we used each investigation. We randomly selected investigations to match the number of aggressive behaviors in the other recordings. As with aggression-triggered SIs and walking-triggered SIs, we compared the number, median latency, and median duration of examples where the investigating male was the social partner to those where the male being investigated was the social partner. When comparing aggregate aggression levels during non-aggressive social behaviors, we used dominant and subordinate identity rather than behavioral state (**S3 Fig K-O**).

#### Random Sampling Procedure

We employed a permutation analysis to ensure that large sample sizes and a subset of examples did not explain the results (**Fig 4A**). Fifty sequences were selected because the recording with the fewest aggression-triggered social interaction sequences had 64 examples. Across all 11 recordings, we calculated the number of sequences in which the aggressor male was the social partner and the number in which the aggressed male was the social partner within the subset of data. We then created an index value using the following formula:

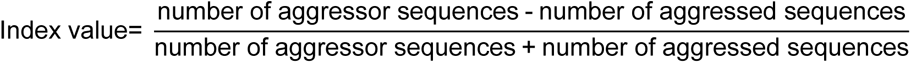

This procedure was repeated 1,000 times to generate a distribution of index values. We repeated this process with the non-social and social control analyses to ensure that our analyses included a representative sample of those behaviors (**Fig 4B-C**). We determined whether the mean index value was significantly different than zero by comparing the data to a normal distribution (mean of 0 and standard deviation of 1) using a z-test.

#### Predicting Behavioral State

We used decision tree classifiers to predict the behavioral state of the male social partner in social interactions following triggering behaviors (aggressive encounters, walks, or investigations), as they can reliably group numerical predictors into two classes with low computational cost (**Fig 3E**) (60). These calculations were performed with the Matlab fitctree function. The predictor variables were the duration of the social interaction and the latency between the behavior and the social interaction. The outcome variable was the behavioral state of the male social partner. For each type of behavioral sequence (aggression-, walking-, and investigation-triggered), we randomly selected 75% of the data as training data. The remaining 25% of the data was used as testing data. The accuracy of the classifier was determined by dividing the total number of correct predictions by the total number of predictions and multiplying by 100 to convert the value to a percentage. Randomly selecting examples to train and test the classifier was repeated 1,000 times. We computed accuracy for each of the 1,000 iterations. Chance levels were 50% because the outcome variable (behavioral state) was a binary value. A z-score was calculated to evaluate whether classifiers performed significantly above chance (α = 0.05). The following formula was used:

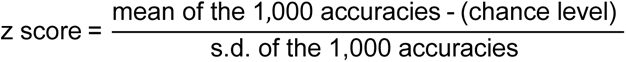

If the z-score was above 1.96, the average accuracy was significantly higher than chance. For all sequence types, we performed this procedure on three sets of data. The first was the observed data, which included every example. The second was a sample-size-matched control dataset, which was used to control for sampling bias. Here, an equal number of aggressed and aggressive examples were used for the training data set. The third was randomized data to disrupt the temporal relationship between aggressive behaviors and social interactions. These procedures were also used to predict the aggregate aggression level of the male social partner (**S3 Fig E**).

#### Predicting Behavioral Sequences

We also used decision tree classifiers to predict the type of behavioral sequence (aggressive or control sequence; **Fig 4D**). This procedure was performed on all observed, size-matched, and randomized data, as described above. Decoding was performed by using the latency, duration, and the behavioral state of the male social partner as predictor variables. The type of sequence was used as the outcome variable. Thus, chance levels were 50%.

#### Temporal Dynamics

To assess the temporal profile of behavioral sequences, we grouped examples into five one-hour bins (**Fig 6**; **S4 Fig**). When assessing the role of behavioral state, we found the proportion of events in each hour where the aggressor/aggressed, walker/non-walker, or investigator/investigated was the male social partner. We also repeated this procedure for aggregate aggression levels. We then trained multi-class support vector machines (mcSVM) to decode the hour in which behavioral sequences occurred using the Matlab fitcecoc function. mcSVMs were used because there were more than two output classes (61). Here, we used the latency between the triggering behavior and the social interaction, the duration of the social interaction, and the identity of the male social partner as predictor variables. The hour in which the sequence began was used as the outcome variable, so chance levels were 20%.

Within each hour, we also asked whether we could predict the identity of the social partner. Decision tree classifiers trained on the latency between the trigger behavior and male-female interaction and the duration of the social interaction were used.

#### Three-step Behavioral Sequences

We characterized three-part behavioral sequences: an aggressive encounter, the social interaction between the aggressed male and a female, and the following social interaction. Based on the male and female involved in the second social interaction, we grouped the sequences into four types (**Fig 7**; **Table 3**). We quantified the proportion of each sequence type across all recordings and the number of each sequence type within each recording.

To assess whether the proportion of each type was different than expected by chance, we used the temporally randomized data (described above). For each sequence type, we generated a random distribution of proportions by calculating the proportion of each type within each shuffle iteration. The actual proportions were compared to the random distributions using a z-score calculated with the following formula:

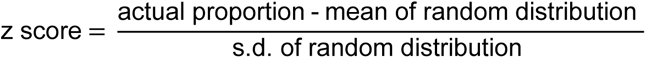

If the z-score was greater than 1.96 or less than 1.96, the actual proportion of events differed significantly from chance.

We also trained a mcSVM to decode the sequence type, given features of the aggressive behavior. We used the average speed during the encounter and the total distance traveled during the behavior for both aggressor and aggressed as predictor variables. The outcome variable was the sequence type (1–4), meaning that chance-level accuracy was 25%. 75% of the data was used as training data, and the remaining 25% was used as testing data. This procedure was performed 1,000 times to generate a distribution of accuracy values, and we computed the z-score using the following formula:

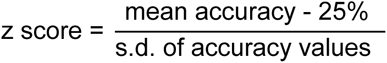

To assess the inter-individual distance between the interacting male and female during the behavioral sequence, we first separated the sequences into four phases: the aggressive encounter, the latency between the aggressive encounter and the social interaction with the aggressed male, the social interaction, and a five second period following the end of the social interaction. We then determined the duration of the longest phase across our entire dataset and used this duration to normalize across all events. For each frame of every sequence, we found the distance between the female social partner and the aggressor and the aggressed males. We then interpolated these distance values to match the length of the longest phase using the Matlab interp function. Thus, all portions of the events were time normalized.

We used a mcSVM to determine the sequence type using inter-individual distances. Predictor variables included the distance between the aggressor and the female at the start of the aggressive behavior, the end of the aggressive behavior, the start of the social interaction, the end of the social interaction, the start of the second social interaction, the end of the second social interaction, and the distances between the aggressed male and the female at the same time points. The outcome variable was sequence type (1, 2, 3, or 4), so chance levels were 25%.

To assess the success of the behavioral strategy, we determined how frequently fights occurred. For each recording, we then calculated how often fights occurred after type 1 or other sequences (2, 3, and 4). Types 2, 3, and 4 were combined because these sequences constituted 47% of the sequences (**Fig 7M**). We also determined the proportion of the total fights that occurred after sequence type 1 and the other sequence types (**Fig 7N**).

## Data and Code Availability

All code and data are available upon request. All data in figures is available as Metadata.

## Author Contributions

Conceptualization – JPN, RSC, MRW

Formal Analysis – RSC

Funding Acquisition – JPN

Data Collection – MRW

Methodology – JPN, RSC, MRW

Project Administration – JPN

Resources – JPN

Software – RSC, MRW, JPN

Validation – RSC, JPN

Data Visualization – RSC, JPN, MRW

Writing – original draft – JPN, RSC

Writing – review & editing – RSC, MRW, JPN

## Acknowledgments

We thank Dr. Lisa Stowers for helpful comments on the manuscript, the staffs of the Life Science Research Facility at the University of Delaware for their assistance in caring for the animals, the High-Performance Computing Group at the University of Delaware for support in maintaining the data processing pipeline, Jim Farmer and Jamie Quesenberry for aid in building lab equipment, and Dan Sangiamo for help collecting the data. This research was funded by NIMH R01MH122752, NIH P20GM103653, the University of Delaware Research Foundation, and Delaware’s General University Research Program.

---

**S1 Fig.**
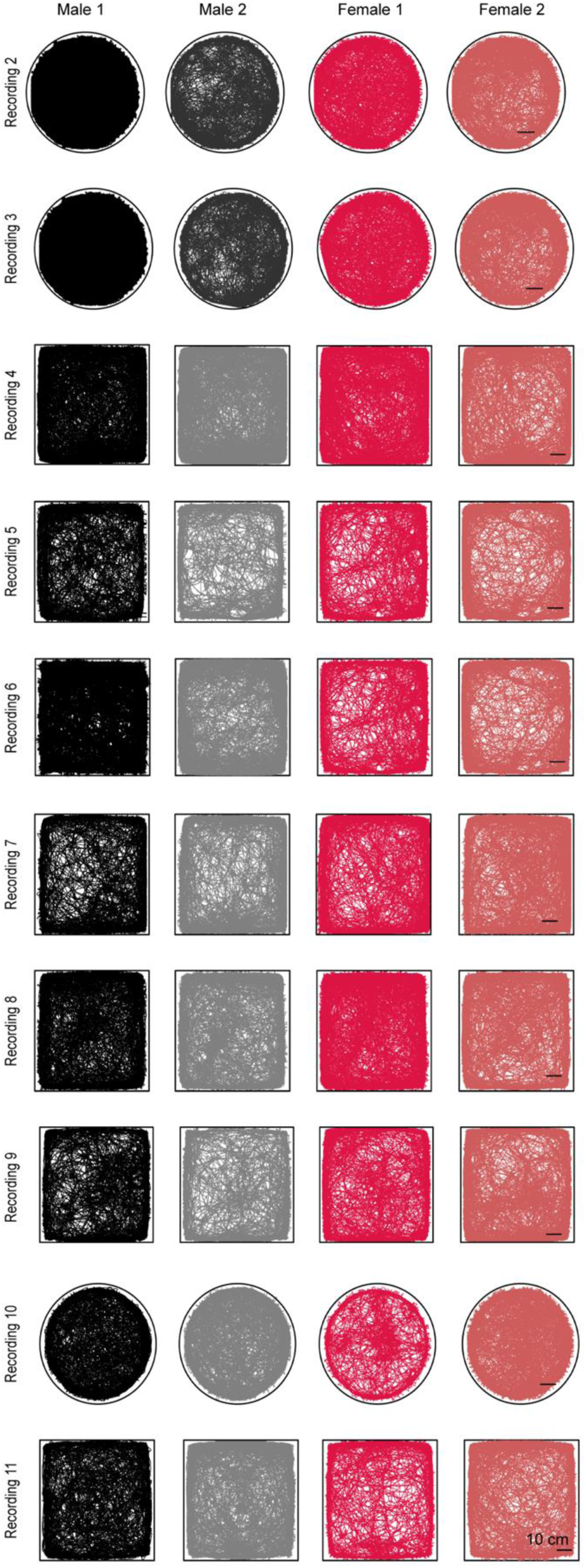
Mouse trajectories. Behavioral trajectories for each mouse across all recordings. Note, all mice explored the majority of the behavioral arena. Data from the first recording was shown in Fig 1C.

**S2 Fig.**
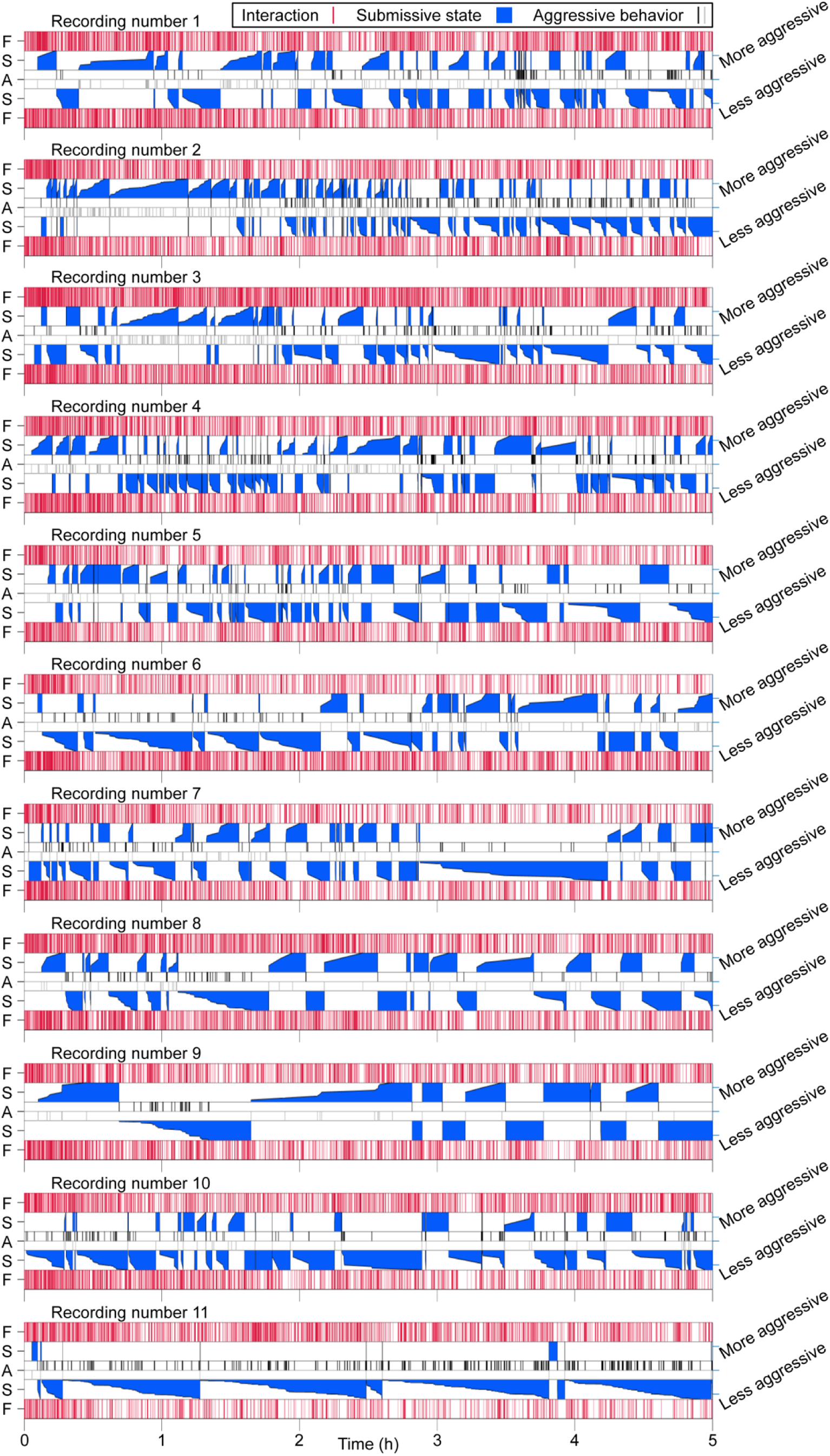
Ethograms depict interactions between mice. The top and bottom rows (red vertical lines, labeled F) show male and female interactions. Middle row denotes aggressive behaviors between males. Top of the central row (black lines, labeled A) shows acts of aggression from the more aggressive animals, while bottom of the central row (gray lines, labeled A) shows acts of aggression from the less aggressive animal. Interleaved rows (labeled S) indicate submissive states (blue patches). Steps in the submissive states indicate consecutive acts of aggression from the other male.

**S3 Fig.**
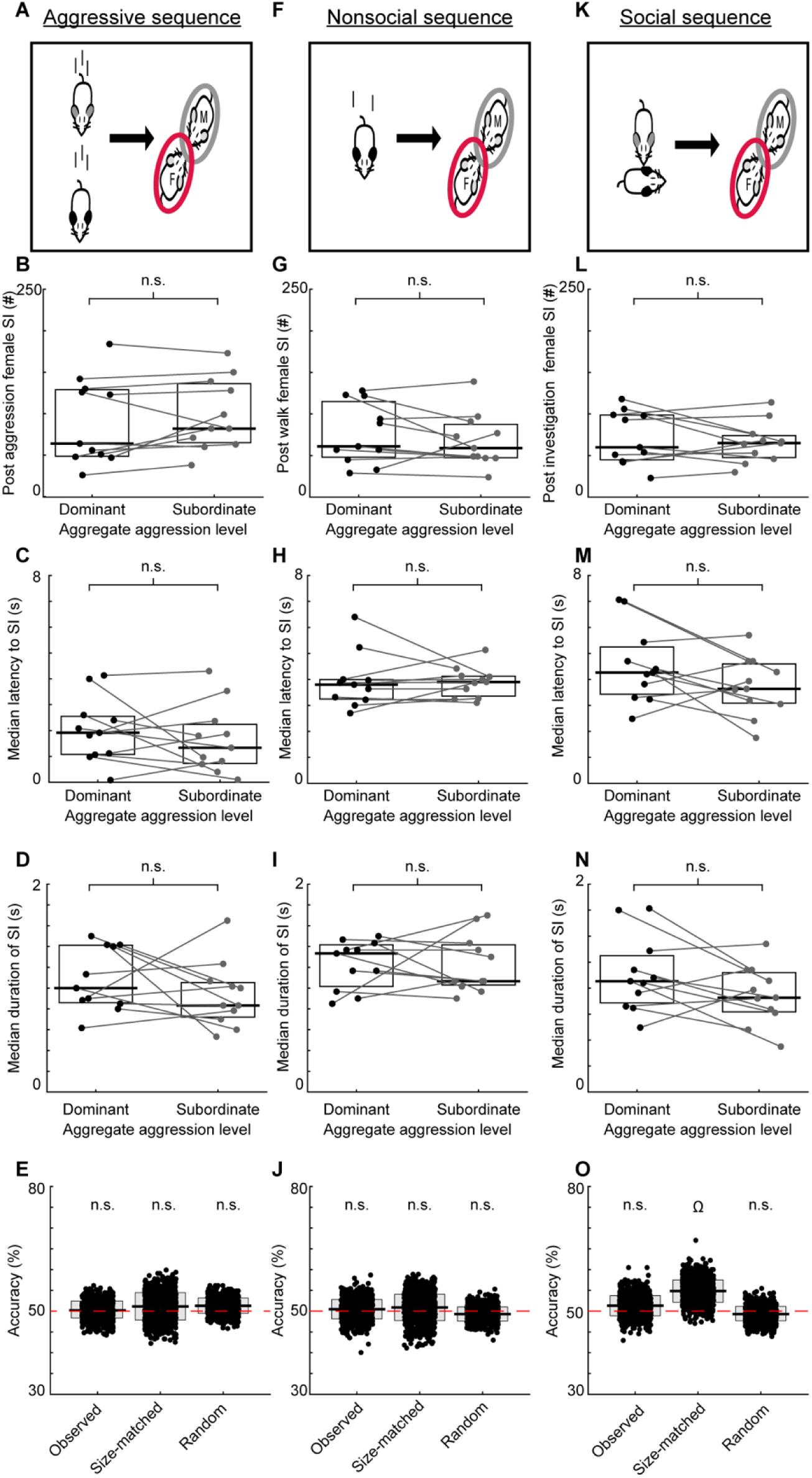
Aggregate aggression levels do not modulate subsequent interactions with females. (A) Schematic of aggressive social sequences. (B) The number of male-female interactions following aggressive behaviors. Lines connect co-recorded mice. The horizontal bars and boxes below the data show the medians and interquartile ranges (25-75%). (C) The latency between aggressive encounters and social interactions. (D) The duration of social interactions following aggressive encounters. (E) Decoders’ performance when predicting the aggregate aggression level of the male social partner in post-aggression social interactions. The horizontal bars and boxes below the data show the means and standard deviations. The red line denotes chance levels. (F) Schematic of nonsocial sequences. Sequences consisted of a male walking in isolation followed by male-female social interactions. (G) As in B, for walking-triggered sequences. (H) As in C, for walking-triggered sequences. (I) As in D, for walking-triggered sequences. (J) As in E, for walking-triggered sequences. (K) Schematic of non-aggressive social sequences. Sequences consisted of investigative male interactions followed by male-female social interactions. (L) As in B, for investigation-triggered sequences. (M) As in C, for investigation-triggered sequences. (N) As in D, for investigation-triggered sequences. (O) As in E, for investigation-triggered sequences. Ω = p < 0.05, n.s. = p ≥ 0.05

**S4 Fig.**
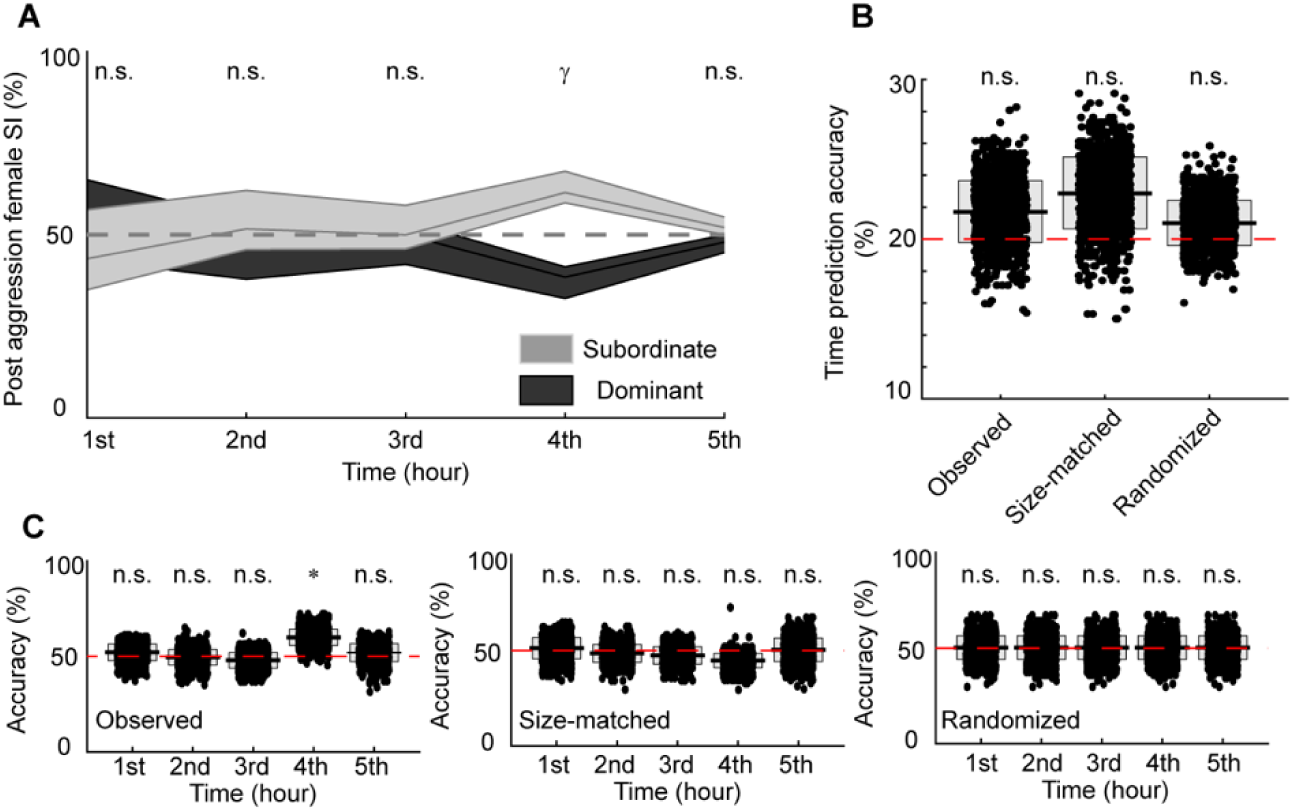
Across time, aggregate aggression values do not modulate subsequent interactions with females. (A) The percentage of aggression-triggered sequences binned hourly for the dominant and subordinate male. The lines and shaded regions show the medians and interquartile ranges (25-75%). (B) Decoders’ performance when predicting when the sequences occurred. The horizontal bars and boxes below the data show the means and standard deviations. The red line denotes chance levels. (C) Decoders’ performance when predicting the behavioral state of the male interacting with the female for all observed data, size-matched controls, and randomized start times of aggressive male-male interactions. * = p < 0.01, γ = p < 0.001, n.s. = p ≥ 0.05

**S1 Table.** Statistical values for analyses presented in S3 Fig. Bold letters indicate corresponding figure panel.

|  |  | Central Tendency/Variability | Statistic | p |
| --- | --- | --- | --- | --- |
| Aggressive Sequences –<br>Number of Instances<br>(**S3-B**) | Dominant | Median/IQR: 64.08 ± 80.25 | W = 17 | 0.17 |
|  | Subordinate | Median/IQR: 82 ± 71.25 |  |  |
| Aggressive Sequences –<br>Median Latency<br>(**S3-C**) | Dominant | Median/IQR: 1.92 ± 1.47 | W = 41.50 | 0.48 |
|  | Subordinate | Median/IQR: 1.33 ± 1.51 |  |  |
| Aggressive Sequences –<br>Median Duration<br>(**S3-D**) | Dominant | Median/IQR: 1 ± 0.55 | W = 44.50 | 0.33 |
|  | Subordinate | Median/IQR: 0.83 ± 0.33 |  |  |
| Aggressive Sequences –<br>Decoder Accuracy<br>Percentage (**S3-E**) | Observed | Mean/STD: 50 ± 2.07 | z = 0.07 | 0.47 |
|  | Size-matched | Mean/STD: 51 ± 3.29 | z = 0.35 | 0.36 |
|  | Randomized | Mean/STD: 51 ± 1.81 | z = 0.71 | 0.24 |
| Non-Social Sequences –<br>Number of Sequences<br>(**S3-G**) | Dominant | Median/IQR: 71 ± 51.25 | W = 53 | 0.08 |
|  | Subordinate | Median/IQR: 55 ± 45.50 |  |  |
| Non-Social Sequences –<br>Median Latency<br>(**S3-H**) | Dominant | Median/IQR: 3.80 ± 0.87 | W = 28 | 0.68 |
|  | Subordinate | Median/IQR: 3.97 ± 0.90 |  |  |
| Non-Social Sequences –<br>Median Duration<br>(**S3-I**) | Dominant | Median/IQR: 1.07 ± 0.28 | W = 9 | 0.05 |
|  | Subordinate | Median/IQR: 1.40 ± 0.35 |  |  |
| Non-Social Sequences –<br>Decoder Accuracy<br>(**S3-J**) | Observed | Mean/STD: 50.45 ± 2.34 | z = 0.19 | 0.42 |
|  | Size-matched | Mean/STD: 50.89 ± 3.18 | z = 0.28 | 0.39 |
|  | Randomized | Mean/STD: 49.31 ± 1.70 | z = -0.41 | 0.34 |
| Social Sequences –<br>Number of Instances<br>(**S3-L**) | Dominant | Median/IQR: 60 ± 30 | W = 31.50 | 0.92 |
|  | Subordinate | Median/IQR: 58 ± 56.75 |  |  |
| Social Sequences –<br>Median Latency<br>(**S3-M**) | Dominant | Median/IQR: 4.67 ± 2.61 | W = 49 | 0.12 |
|  | Subordinate | Median/IQR: 3.08 ± 2.59 |  |  |
| Social Sequences –<br>Median Duration<br>(**S3-N**) | Dominant | Median/IQR: 1.07 ± 0.22 | W = 37 | 0.75 |
|  | Subordinate | Median/IQR: 1 ± 0.20 |  |  |
| Social Sequences –<br>Decoder Accuracy<br>(**S3-O**) | Observed | Mean/STD: 51.31 ± 2.34 | z = 0.54 | 0.29 |
|  | Size-matched | Mean/STD: 54.83 ± 2.70 | z = 1.79 | 0.04 |
|  | Randomized | Mean/STD: 49.31 ± 1.70 | z = -0.32 | 0.35 |

**S2 Table.** Statistical results corresponding to analyses presented in S4 Fig. Bold letters indicate corresponding figure panel.

|  |  | Central Tendency/Variability | Statistic | p |
| --- | --- | --- | --- | --- |
| Percentage of Examples: Dominant/Subordinate (**S4-A**) | Hour 1 | Median/IQR: 56.25 ± 21.84/43.48 ± 21.84 | W = 41 | 0.52 |
|  | Hour 2 | Median/IQR: 48.39 ± 16.09/51.61 ± 16.09 | W = 26 | 0.58 |
|  | Hour 3 | Median/IQR: 50 ± 11.89/50 ± 11.80 | W = 25.50 | 0.87 |
|  | Hour 4 | Median/IQR: 48.46 ± 8.59/61.54 ± 8.59 | W = 1 | < 0.001 |
|  | Hour 5 | Median/IQR: 48 ± 4.84/52 ± 4.84 | W = 16 | 0.50 |
| Decoder Accuracy Percentage (**S4-B**) | Observed | Mean/STD: 21.71 ± 1.95 | z = 0.87 | 0.19 |
|  | Size-matched | Mean/STD: 22.87 ± 2.28 | z = 1.26 | 0.10 |
|  | Randomized | Mean/STD: 21 ± 1.43 | z = 0.70 | 0.24 |
| Decoder Accuracy Percentage: Observed (**S4-C**) | Hour 1 | Mean/STD: 52.08 ± 4.35 | z = 0.48 | 0.23 |
|  | Hour 2 | Mean/STD: 48.54 ± 4.04 | z = -0.11 | 0.46 |
|  | Hour 3 | Mean/STD: 47.91 ± 4.71 | z = -0.50 | 0.41 |
|  | Hour 4 | Mean/STD: 59.97 ± 4.26 | z = 2.34 | 0.01 |
|  | Hour 5 | Mean/STD: 51.93 ± 4.66 | z = 0.41 | 0.34 |
| Decoder Accuracy Percentage: Size-matched (**S4-C**) | Hour 1 | Mean/STD: 51.12 ± 5.61 | z = 0.20 | 0.42 |
|  | Hour 2 | Mean/STD: 48.41 ± 4.68 | z = -0.34 | 0.37 |
|  | Hour 3 | Mean/STD: 47.34 ± 4.50 | z = -0.59 | 0.28 |
|  | Hour 4 | Mean/STD: 44.63 ± 3.74 | z = -1.43 | 0.08 |
|  | Hour 5 | Mean/STD: 50.25 ± 6.19 | z = 0.04 | 0.48 |
| Decoder Accuracy Percentage: Randomized (**S4-C**) | Hour 1 | Mean/STD: 52.55 ± 3.85 | z = 0.66 | 0.25 |
|  | Hour 2 | Mean/STD: 49.94 ± 3.94 | z = -0.02 | 0.49 |
|  | Hour 3 | Mean/STD: 48.36 ± 3.90 | z = -0.42 | 0.34 |
|  | Hour 4 | Mean/STD: 49.97 ± 3.91 | z = -0.01 | 0.50 |
|  | Hour 5 | Mean/STD: 48.73 ± 3.75 | z = -0.34 | 0.37 |

## Notes

### Competing Interest Statement

The authors have declared no competing interest.

### Summary of Updates

Updated manuscript includes the supporting information.

